# Adaptive molecular convergence is pervasive across deep time and largely decoupled from phenotypic convergence

**DOI:** 10.64898/2026.04.23.718300

**Authors:** Cory A. Berger, Marina I. Stoilova, Rebecca M. Varney, Sam C. Abrams, Maria-Pia Miglietta, Paulyn Cartwright, Todd H. Oakley

## Abstract

Researchers often infer evolutionary repeatability when selection scans implicate homologous genes in repeatedly evolved traits or ecologies. However, the causes and frequency of genome-scale molecular convergence remain unresolved, particularly over deep time. We show that adaptive molecular convergence— excess convergence of nonsynonymous substitutions, consistent with positive selection—is pervasive across Medusozoa. Molecular convergence declines over time but persists among lineages separated by *>* 600 million years, exceeding null expectations based on random overlap. However, lineages sharing repeatedly evolved phenotypes (eyes, medusa loss, upright colonies) do not exhibit elevated molecular convergence relative to other comparisons. Instead, convergence is non-randomly distributed across genes and enriched for environment-facing functions, including metabolism, immunity, and xenobiotic processing, suggesting that widespread reuse of genes reflects multifaceted organism-environment interactions.

---

A central question in biology is how often similar ecological challenges lead to repeated genetic changes, and how reliably those changes explain repeated phenotypic adaptations. Repeated origins of similar phenotypes across independent lineages suggest that natural selection drives evolution towards similar solutions under comparable ecological conditions. Furthermore, the use of homologous genes in convergent origins of similar traits or ecological contexts suggests that genetic paths may also be repeatable (*1–4*). However, establishing causal links between genes and phenotypes remains difficult because we lack functional information for most genes in most species (*5*). Consequently, biologists often rely on comparative genomic approaches that use signatures of selection to associate genes with repeated adaptations. These genome-wide approaches implicitly assume that the molecular convergence observed between phenotypically convergent taxa reflects the genetic basis of observed adaptations. However, adaptive molecular convergence may also reflect broader patterns of selection acting across many genes and ecological contexts, rather than the genetics of any particular trait (*6*). Distinguishing these possibilities requires understanding how frequently adaptive molecular convergence occurs across genomes and over deep evolutionary timescales, and whether it is systematically associated with particular phenotypic adaptations.

Patterns of molecular convergence across evolutionary time have already provided important perspectives, especially over relatively shallow timescales. An emerging pattern is a negative relationship between gene reuse and divergence time (*2, 7*), suggesting that molecular convergence between distant lineages may be rare. However, the magnitude and rate of this decline are poorly understood across deep timescales (> 100 My), as we have little data concerning the frequency of molecular convergence between such distantly related species. It is also unclear whether this decline reflects decreasing reuse of genes underlying particular convergent adaptations or a broader decline in molecular convergence without direct relationships to phenotype and ecology. It is challenging to draw general conclusions from the literature due to extensive heterogeneity in the datasets and statistical approaches used across studies and historical difficulties differentiating between adaptive and neutral molecular convergence (*8,9*). In addition, many analyses focus on single-copy orthologs, which represent a small and potentially biased subset of genes (*10, 11*). Recent methodological advances help overcome these limitations. For example, CSUBST detects convergent amino acid substitutions consistent with positive selection across homologous proteins and can be applied to large multi-copy gene families without focusing only on predefined convergent lineages (*12*). When combined with new phylogenetic methods for analyzing lineage-pair data—quantities defined for pairs of taxa such as convergence and divergence time (*13*)—these approaches allow genome-wide comparisons of adaptive molecular convergence across lineages and timescales within a unified statistical framework.

Here, we examine patterns of adaptive molecular convergence across Medusozoa (jellyfish and hydroids), an ancient and diverse clade in which key ecological phenotypes have evolved repeatedly over deep evolutionary time. These include multiple origins of eyes (*14*), repeated losses of the free-swimming medusa stage (*15*), and transitions in colony architecture (*16*). Because these convergent traits span hundreds of millions of years and diverse life histories, Medusozoa provides an unusually powerful system to test whether genome-wide molecular convergence predictably tracks repeated phenotypic adaptation over deep time. We quantify adaptive molecular convergence across thousands of gene families and compare its distribution across lineages differing in phenotype, divergence time, and ecological context. Our results reveal widespread molecular convergence that is not predictably associated with repeated phenotypic evolution, suggesting that protein sequence evolution can be strikingly repeatable even between distant relatives, while the selective causes of that repeatability are often diffuse, lineage-specific, and not easily identifiable from phenotype alone.

## Medusozoa originated more than 650 Mya and underwent extensive convergent evolution of phenotypes

To better understand the timing and patterns of repeated evolution in Medusozoa, we used new and existing transcriptomic and genomic datasets for phylogenetic and character evolution analyses. Bayesian relaxed molecular clock analyses using 17 fossil calibrations (data S3) firmly support a Cryogenian origin of Medusozoa (c. 685 Mya) and early diversification of major extant clades, including Ediacaran origins of Hydrozoa and Scyphozoa, an origin of Cubozoa near the early Cambrian, and Staurozoa later in the Paleozoic (Fig. 1). The resulting phylogeny resolves several previously uncertain relationships, most notably the placements of Capitata, Leptothecata, and the polyphyletic Filifera (Fig. 1; figs. S10, S11).

**Figure 1:**
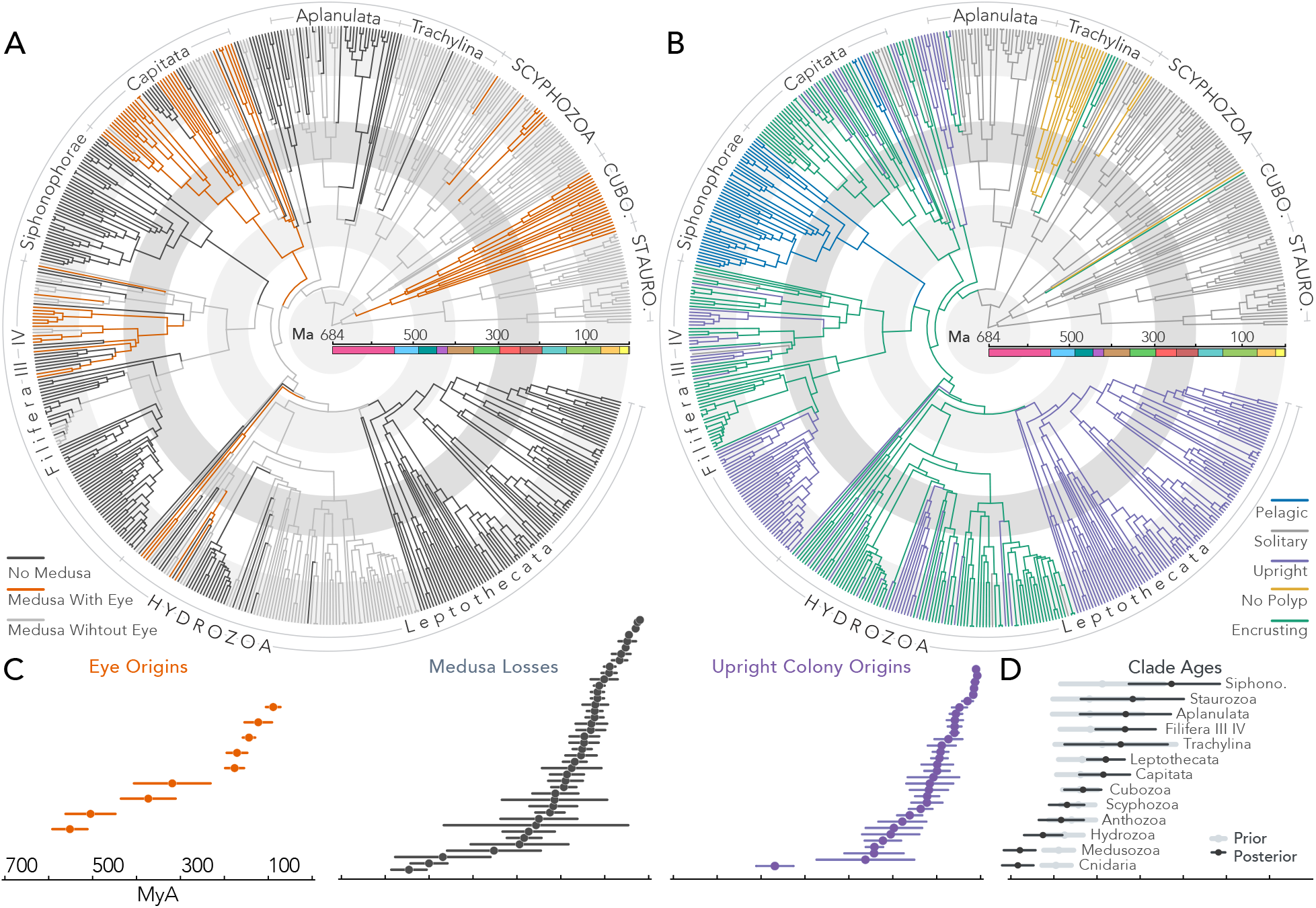
Convergent character transitions have occurred dozens of times across 680 My of medusozoan evolution. (A) Joint ancestral state reconstructions (ASR) of medusa stage and eye presence. ASRs were inferred using corHMM under the best-fit model by AIC (data S4). Time-calibrated phylogeny was inferred from an MCMCtree analysis of 68 taxa chosen to evenly sample major clades; these times were transferred to a 561-taxon phylogeny for ASR. Filifera I-II are not highlighted here for space but can be found in the full trees on Dryad. “No medusa” includes medusoids, sporosacs, and absent gonophores. Medusa and eye presence were plotted together because these characters are correlated (eyes only occur on medusae; see Methods). Full species trees with taxon labels and branch support values can be found on the Dryad and Github; see fig. S16 for ASR of eyes only and fig. S17 for ASR of medusa stage with detailed character categories. (B) ASR of colony architecture. (C) Ages of key character transitions. Points and error bars show the mean and 95% CI of node ages calculated from 500 chronograms sampled from the MCMCtree posterior distribution. (D) Estimated ages of key clades. Points and error bars show the mean and 95% highest-posterior density intervals for the prior (fossil calibrations without sequence data) and posterior.

Using this time-calibrated phylogenetic framework, we reconstructed the evolutionary histories of multiple traits across Medusozoa, revealing extensive convergent evolution over deep time. At least nine independent origins of eyes, 41 losses of the medusa stage, and 32 transitions from encrusting to upright colony architectures were inferred (Fig. 1C). These repeatedly evolved traits are associated with distinct life histories and ecological strategies (*15, 17, 18*), providing a natural framework for testing links between phenotypic and molecular convergence. Across all shree traits, the divergence times between lineages exhibiting convergent phenotypes spanned approximately 200–600 million years, enabling comparisons between molecular and phenotypic convergence across deep evolutionary timescales.

### Adaptive molecular convergence exceeds null expectations across deep time

We quantified adaptive molecular convergence, defined as an excess of convergent nonsynonymous substitutions consistent with positive selection, across 86 medusozoan species using CSUBST (*12*). CSUBST identifies branch pairs in gene trees with an elevated rate of convergent nonsynonymous substitutions compared to the rate of convergent synonymous substitutions. This ratio, *ω*_*c*_, is analogous to but distinct from the well-known measure of sequence evolution *ω*; the neutral expectation of *ω*_*c*_ is 1, and *ω*_*c*_ ≫ 1 is evidence of convergent selection towards identical amino acids. For each pair of species, we calculated molecular convergence as the proportion of gene families with at least one convergent branch pair, not distinguishing between orthologs and paralogs (see Methods).

Adaptive molecular convergence occurred broadly across medusozoan lineages and genes. In total, we identified 426,322 convergent branch pairs (0.091% of all tested branch pairs) across 8991 of 9194 gene families. Phylogenetic regression showed that the degree of molecular convergence declined significantly with increasing divergence time (Fig. 2A; *p* ≪ 0.01), consistent with predictions that distantly-related taxa have fewer opportunities for shared substitutions (*2*). Nonetheless, there was substantial variation across all divergence times, indicating that phylogenetic relatedness and evolutionary time explained only a small fraction of the extensive molecular convergence in our data (residual standard error [RSE] = 0.82 on the transformed scale). Sequence data simulated from our alignments under a scenario of zero convergence resulted in a low false positive rate and no relationship with divergence time (fig. S6; Supplementary Text), indicating that observed patterns of convergence cannot be attributed to biases in the underlying alignments and gene trees. Furthermore, convergent substitutions identified by CSUBST were less physicochemically conservative than other substitutions in the same proteins (fig. S7). This pattern was not observed in the data simulated without convergent selection, bolstering inferences that most convergence events identified by CSUBST were driven by positive selection.

**Figure 2:**
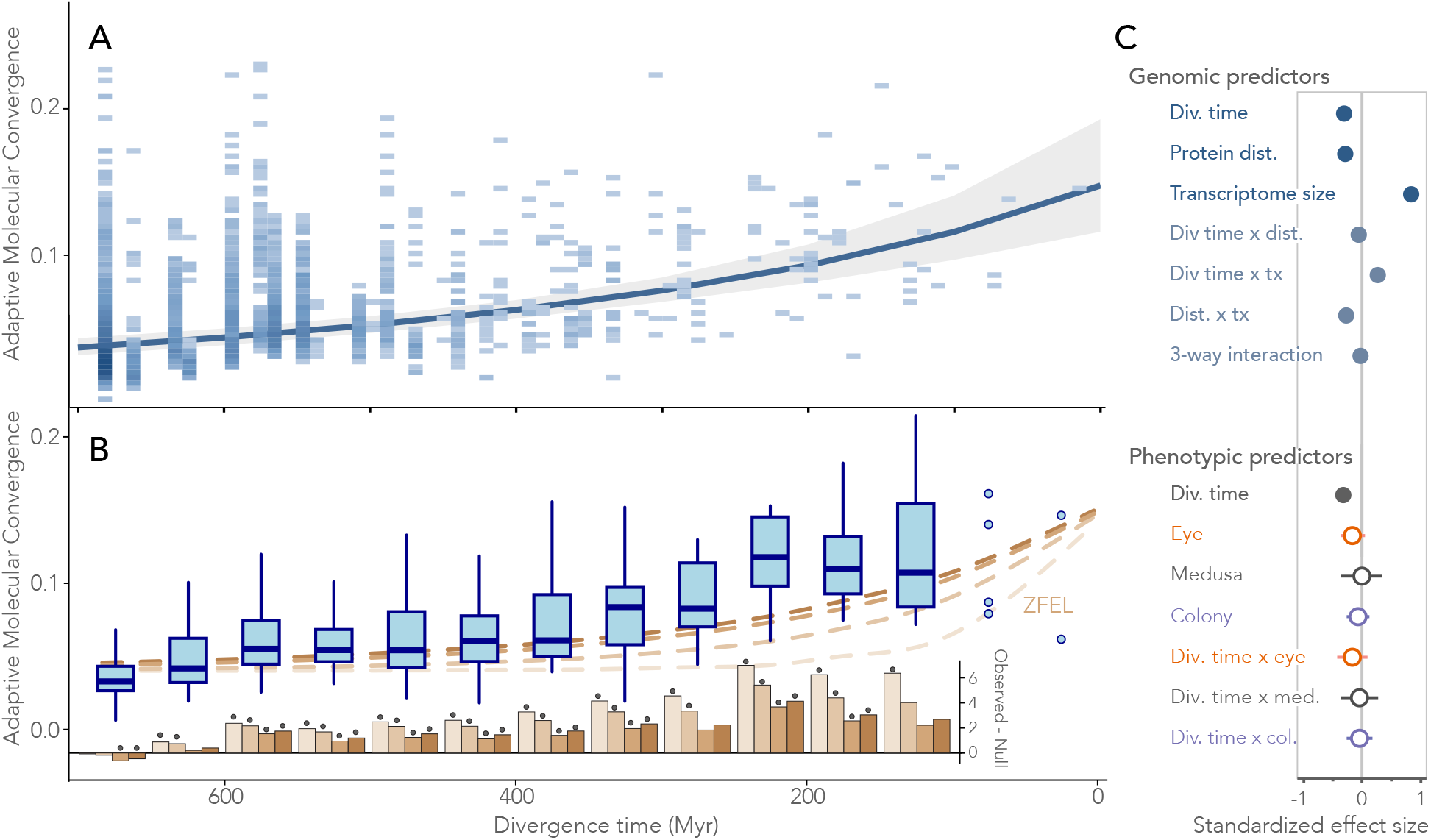
Adaptive molecular convergence declines with divergence time and exceeds null expectations. (A) Adaptive molecular convergence (AMC) among medusozoan lineages as a function of divergence time. AMC was quantified as the proportion of gene families exhibiting convergent branch pairs. AMC values were transformed with a Yeo-Johnson transformation (λ = −0.565); the regression is plotted here back-transformed onto the original scale. Points representing pairwise species comparisons (*n* = 3570) were binned for visualization purposes, with darker shading indicating higher observation density. The solid line shows the phylogenetic regression between AMC and divergence time, with the shaded ribbon indicating the model confidence interval. (B) Observed AMC compared with expectations under Zero-Force Evolutionary Law (ZFEL) simulations. Boxplots show the distribution of observed convergence (data in Panel A) binned into 50 My intervals. Individual points were plotted for intervals with < 10 data points. Dashed lines show median convergence expected from ZFEL simulations, which model the decline in similarity among independently evolving systems without correlated selection. The multiple lines show different realizations of the null with varying rates of loss. All scenarios are set to converge on the same steady-state value, which equals the convergence in the final time point of observed data (∼ 4%; see Methods). Below, bars show mean differences between observed and null values within each bin. Asterisks indicate *p* < 0.05 (Bonferroni-corrected within each null model). The most conservative null distribution (dark brown) was constructed such that its lower 0.025 quantile would include the observed value at our oldest time point; this explains why null values slightly exceed observed values in the oldest bin (see fig. S14 and Supplementary Text). (C) Predictors of adaptive molecular convergence. Top, standardized regression coefficients from a model evaluating the effects of divergence time, protein sequence distance, transcriptome size, and their interactions on AMC. Confidence intervals are smaller than points. Bottom, standardized regression coefficients for a model testing whether eyes, medusa stage, and colony form predict AMC. Horizontal bars indicate confidence intervals and shaded circles indicate significance (*p* < 0.05).

Do levels of adaptive molecular convergence across Medusozoa reflect systematic constraints and selection, or can they simply arise from random overlaps among loci in finite genomes? To test this, we simulated null expectations of convergence derived from the Zero-Force Evolutionary Law (*19*), which states that similarity among independently evolving systems declines over time in the absence of constraints or directional forces, such as selection acting the same way in both systems. Accordingly, if molecular convergence reflects only uncorrelated selection across genes, it will decline with increasing divergence time and not exceed the levels expected from random overlap alone (see Methods).

Levels of adaptive molecular convergence in Medusozoa consistently exceeded these null expectations, even across deep evolutionary time. Across a 650 My range of divergence, molecular convergence was significantly higher than predicted under random overlap alone (Fig. 2B). Null models were constructed under a range of possible parameter values (see Methods), and convergence exceeded null predictions across most time intervals even under the most conservative conditions (dark brown in Fig. 2B), which correspond to the slowest possible decline compatible with uncorrelated evolution. The persistent excess of observed convergence under these very conservative conditions indicates that adaptive molecular convergence in Medusozoa reflects systematic evolutionary forces and/or constraints operating across deep time.

### Adaptive molecular convergence is not explained by phenotypic convergence

If adaptive molecular convergence primarily reflects reuse of genes underlying particular convergent traits, then species with those traits should show higher levels of molecular convergence. To test this prediction, we compared levels of molecular convergence among species pairs with phenotypic convergence of eyes, upright colony architectures, or losses of the medusa stage to those of other species pairs without obvious convergent traits. Contrary to expectations, species with convergent phenotypes were statistically indistinguishable from other taxa in both overall levels of molecular convergence (*p* > 0.1) and the effect of divergence time (*p* > 0.1) (Fig. 2C). Therefore, these three prominent, ecologically important, and repeatedly evolved phenotypes do not explain the observed genome-wide patterns of adaptive molecular convergence.

We next asked whether other biological factors could account for variation in convergence among species pairs. Incorporating pairwise protein distance and transcriptome size as covariates in addition to divergence time improved regression model fit, and all main effects and interactions were significant (Fig. 2C; table S5). As expected, species with greater protein sequence similarity exhibited higher overall levels of molecular convergence and maintained elevated convergence over longer evolutionary intervals. The same was true for species with larger transcriptomes, likely because additional gene copies provide more opportunities for convergence within gene families. This suggests that gene duplication may be an important contributor to patterns of molecular convergence by providing evolutionary substrates for convergent selection. Nevertheless, substantial unexplained variation remained even after accounting for these covariates (RSE = 0.77). We further tested whether adaptive molecular convergence was systematically associated with particular clades or taxonomic groups. After accounting for transcriptome size, species-averaged convergence values lacked detectable phylogenetic signal (*p* = 0.8), and taxa with high or low convergence showed no obvious shared characteristics (fig. S8).

Together, these results indicate that adaptive molecular convergence varies widely among species pairs and cannot be readily attributed to phenotypic similarity or phylogenetic structure. Consistent with this decoupling, although genes with established roles in eye development were moderately enriched in comparisons between eye-bearing species, these genes also often experienced convergent selection in species lacking eyes (fig. S15, table S2). Thus, genomic scans may enrich for trait-related genes, but the pervasive background of adaptive molecular convergence makes it difficult to interpret apparent gene reuse as evidence of trait-specific adaptation.

### Adaptive molecular convergence reflects multi-faceted organism-environment interactions

Convergence may exceed null expectations and be unequally distributed across gene families due to either shared genomic constraints [e.g., similar mutational landscapes, genome architectures, and gene-regulatory networks; (*1, 20*)] or shared ecological contexts resulting in similar adaptive landscapes (*21*). We can partially disentangle these factors because genomic constraints should decline with phylogenetic distance (*22–25*), whereas ecological distance may lack phylogenetic signal (*26, 27*), particularly across divergent clades. We therefore predict that, if convergence is primarily structured by genomic constraints, then closely-related taxa will share more similar sets of convergent genes.

To test this, we quantified the frequency with which members of the same gene family independently experienced convergent selection across different species pairs using an extension of Yeaman et al.’s C-score metric (*28*). Convergent gene families were similar across species quartets (pairs of pairs) more frequently than expected under random chance (mean C-score = 4.91 ± 2.72), with 33.9% of quartets showing significant overlap (Bonferroni-corrected *p* < 0.01). However, similarity in the identities of convergently evolving genes showed no meaningful relationship with phylogenetic distance (*r*^2^ = 0.0011; Fig. 3A). Thus, although adaptive molecular convergence is repeatable at the gene-family level, it is not preferentially shared among closely related taxa, arguing against a dominant role for shared genomic constraints in structuring molecular convergence across Medusozoa. Patterns of molecular convergence were similarly unstructured at the level of gene function. Gene ontology (GO) terms enriched among convergent genes were unevenly distributed across species pairs, with most terms restricted to one or two comparisons (Gini coefficient = 0.78; Fig. 3A). Semantic similarity of enriched GO terms among species quartets showed no relationship with phylogenetic distance (*r*^2^ = 1 × 10^−5^, Fig. 3B), indicating that convergently adapted functions are largely lineage-specific and lack phylogenetic structure. These results further suggest that genomic constraints alone cannot explain which genes or functions repeatedly experience convergent selection.

**Figure 3:**
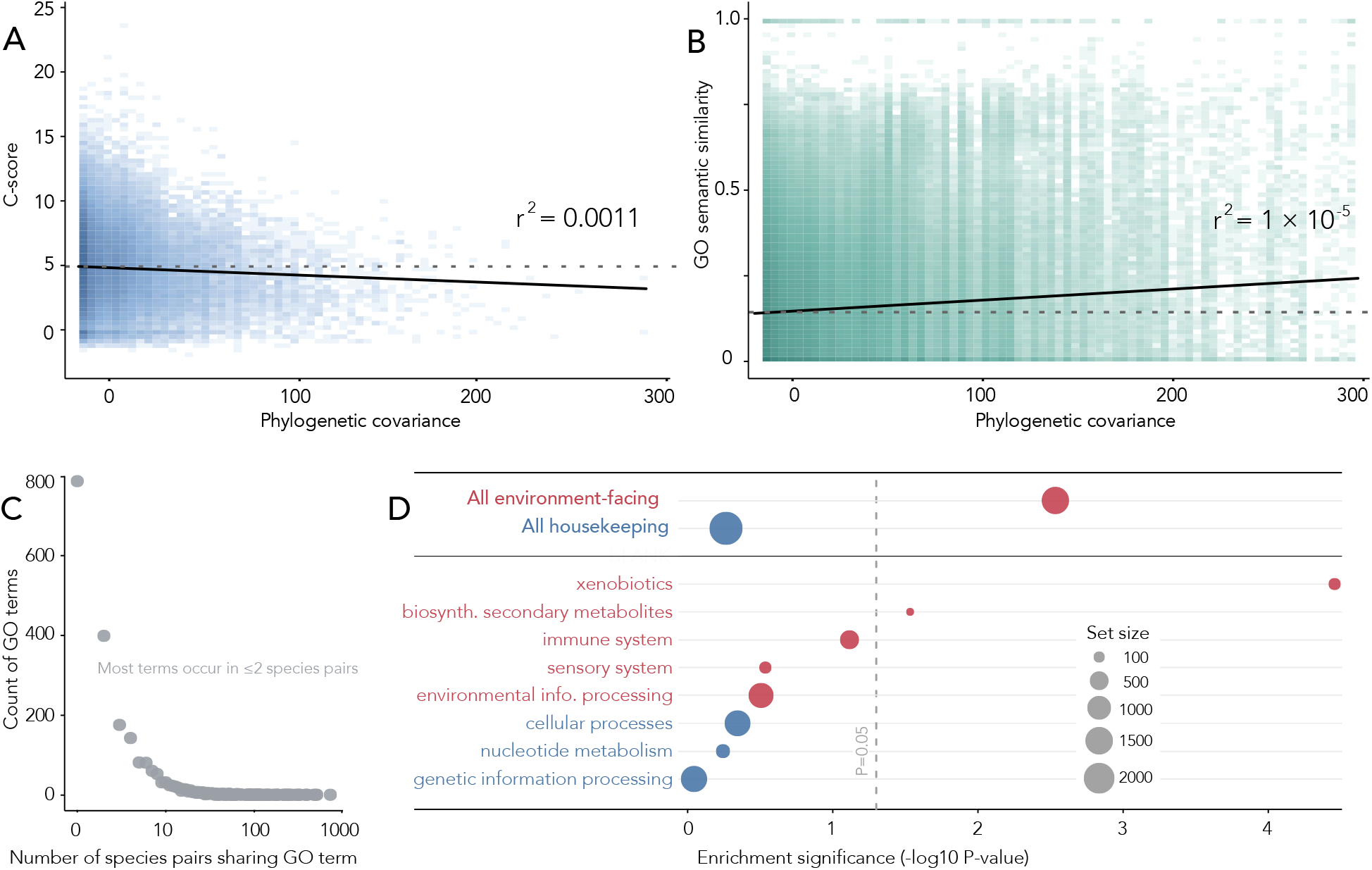
Adaptive molecular convergence lacks phylogenetic structure but is associated with environment-facing genes. (A) Identities of convergent genes are not associated with phylogenetic distance. C-scores represent the overlap of gene families that independently experienced convergence in two pairs of species, with each unit increase representing one standard deviation above the expected overlap from a null hypergeometric distribution. Each point shows the C-score of a randomly-chosen species quartet (n=236,070) plotted against the phylogenetic covariance of that quartet, with darker shading indicating higher observation density. Higher covariance values indicate more shared phylogenetic history. Dashed line indicates average value and solid line indicates a linear regression. (B) GO annotations of convergent genes are not associated with phylogenetic distance. Each point represents the mean semantic similarity of the lists of GO terms enriched among convergent genes between two pairs of species (n=4,755,188 quartets) plotted against phylogenetic covariance. Semantic similarities were calculated as the mean of the maximum similarity between each GO term and members of the other list. Dashed line indicates average value and solid line indicates a linear regression. (C) Number of species pairs with enrichment of each GO term (log scale on X axis). Most GO terms are only found in one or two species pairs. (D) Gene set enrichment analysis of “environment-facing” and “housekeeping” genes. Genes were categorized based on KEGG annotations and grouped into the sub-categories shown here and in table S4. Dashed line indicates raw p-value = 0.05, and point sizes indicate the number of annotated genes in that category.

Despite this lack of phylogenetic structure, adaptive molecular convergence was strongly non-random with respect to gene function across Medusozoa. Gene families with high convergence rates were enriched for functions related to energy metabolism, immune response, and xenobiotic processing, whereas genes with low convergence rates were enriched for internal regulatory processes, including gene regulation, meiosis, and the cell cycle (fig. S9). Using an alternative classification based on KEGG pathway annotations, we grouped genes *a priori* into environment-facing and housekeeping categories and tested for differences in convergence between these classes (table S4). Environment-facing genes were significantly enriched among convergent genes (*q* = 0.00308; Fig. 3D), whereas housekeeping genes were not (*q* = 0.286). Together, these results indicate that adaptive molecular convergence is concentrated in genes involved in diverse organism-environment interactions rather than internal housekeeping functions, consistent with selection associated with multiple ecological axes across deep evolutionary time.

## Discussion

We find adaptive molecular convergence to be pervasive across medusozoan evolution over deep time. However, this convergence is not predictably associated with phenotypic or phylogenetic similarity, and instead reflects broader, lineage-specific ecological pressures. Our results suggest that adaptive molecular convergence is widespread but usually difficult to relate to specific phenotypes or selective agents. Here, we discuss the implications of this pattern for three related questions: the drivers of molecular convergence over deep time, the interpretation of genomic signatures of adaptation, and the predictability of evolutionary outcomes.

First, we argue that molecular convergence over deep time is not primarily structured by shared genomic constraints, but instead reflects selection across complex ecological dimensions. The identities and functions of convergent genes are not associated with phylogenetic distance in Medusozoa, indicating that constraints, which should decay with genetic divergence (*22–25*), do not determine which genes repeatedly experience convergent selection. We do not exclude that constraints influence molecular convergence (*1,29*), but our data suggest that genome-wide patterns of adaptive convergence are mainly driven by selection on diverse, often unobserved, ecological factors. As a result, genes that interface with the external environment show elevated rates of convergence overall, and individual events of molecular convergence occur idiosyncratically across lineages. This pattern is consistent with highly multidimensional selective regimes (*6, 30*). Species therefore experience many uncorrelated and lineage-specific selective pressures that may overlap among taxa, such that the underlying selective agents are often unknown.

Second, this perspective reshapes interpretations of genome-wide molecular convergence and helps reconcile seemingly incongruent patterns in the literature. On the one hand, many studies have identified repeated use of specific genes or pathways in association with convergent phenotypes or ecological transitions (*2, 11*). On the other hand, genome-wide analyses have often failed to detect elevated molecular convergence in lineages with repeatedly evolved traits (*31–36*), a pattern frequently attributed to a pervasive background of neutral convergence (*4, 37*); but see (*33, 34*). We show that, in addition to a neutral background (*9*), there is also a widespread background of adaptive convergence that is not restricted to specific traits. As a result, molecular convergence is not necessarily enriched in focal lineages, even when phenotypes have evolved repeatedly, because trait-specific signals are diluted within this adaptive background. This perspective also reframes patterns such as the reported decline in gene reuse over time (*2*). Although the decline is real, it may not reflect decreasing reuse of genes underlying specific adaptations. Our results suggest that a similar pattern arises even from comparisons among randomly chosen taxa. An important direction for future studies will be to explicitly quantify axes of ecological variation, which could account for some of the variables driving convergent selection (*21, 30*).

We used amino acid substitutions as a signature of convergent selection because this approach is uniquely tractable across long timescales and large gene families. However, further work is needed to understand patterns of convergence across time at other genetic and molecular levels, which can be uncoupled from one another (*38*). Regulatory evolution, for instance, may be more important than protein sequence changes for many convergent phenotypes, particularly morphological traits (*39, 40*). Therefore, integrating multiple signatures of convergence across coding sequences, regulatory regions, and gene expression may provide a more complete view of the genomic basis of repeated adaptation (*41*). However, such analyses remain limited in taxonomic and temporal scope by the availability of high-quality genomes, and further theoretical advances are also needed to better associate regulatory and expression changes with positive selection [e.g., (*42*)].

Finally, our results reveal a tension between two perspectives on evolutionary repeatability. The pervasive nature of adaptive molecular convergence indicates that protein evolution is unexpectedly repeatable even among highly divergent lineages. At the same time, the lack of consistent mapping between molecular convergence and phenotype suggests that this repeatability is not easily interpretable or predictive. This is highlighted by the observation that homologs of known eye-related genes consistently experienced convergent selection in species without eyes. More generally, genomic signatures of selection do not directly identify the ecological causes of selection, and processes such as pleiotropy, epistasis, and functional divergence can produce convergent substitutions with distinct causes and consequences. Similar principles apply at other levels of organization, including morphology: similar phenotypes can arise from different selective pressures and similar selective pressures can produce different phenotypes (*38*). Overall, ecology appears to broadly shape patterns of convergent evolution across deep time, but linking genetic changes to specific traits or selective regimes will require a deeper understanding of genotype–phenotype relationships.

## Acknowledgments

We would like to thank F. Diskin for helping with the medusa morphology literature search; J. Wolfe, P. Nosil, K. Peichel, J. Feder, Z. Gompert, members of the Oakley lab, and the EEMB Evolution discussion group for insightful remarks on the manuscript; A. Collins for providing *Scolionema* samples; and the Smithsonian Tropical Research Institute (STRI), Bocas del Toro Research Station (BRS), R. Collin, and members of the Benthic Cnidaria class at BRS for assistance with sample collection.

## Author contributions

Conceptualization: C.A.B., M.M., P.C., T.H.O; Resources: R.M.V., M.M., P.C.; Investigation: C.A.B., M.I.S., R.M.V., S.C.A; Formal analysis: C.A.B, S.C.A.; Visualization: C.A.B., T.H.O.; Funding acquisition: M.M., P.C., T.H.O.; Supervision: M.M., P.C., T.H.O.; Writing—original draft: C.A.B., T.H.O.; Writing—review & editing: all authors.

## Data and materials availability

Sequence data generated for this study are available at NCBI under Bioproject PRJNA1455705, and accessions of previously published data can be found in Data S1. Transcriptome assemblies, alignments, gene and species trees, CSUBST output, and other large data files will be available on Dryad upon publication. Scripts and smaller data files are available at https://github.com/ucsb-oakley-lab/Medusozoa_project and will also be archived on Dryad.

## Supplementary materials

Materials and Methods

Supplementary Text

Figs. S1 to S18

Tables S1 to S6

References (*43-84*)

Data S1 to S5

