## Supplementary material for "Adaptive molecular convergence is pervasive across deep time and largely decoupled from phenotypic convergence": Methods and Supplemental Material

**This PDF file includes:**

Materials and Methods

Supplementary Text

Figures S1 to S18

Tables S1 to S6

Captions for Data S1 to S6

**Other Supplementary Materials for this manuscript:**

Data S1 to S6

### Materials and Methods

#### Assembly construction and phylogenetics

We assembled a dataset of 86 medusozoan genomes and transcriptomes and 12 anthozoan outgroups, chosen to sample all major medusozoan and anthozoan clades; 22 transcriptomes were newly generated for this study (Data S1). The only group that we did not sample was Endocnidozoa, because these tiny endoparasites are notoriously challenging for phylogenetic analyses due to prevalent host contamination, extensive gene loss, and rapid sequence divergence. Phylogenomic studies place Endocnidozoa sister to the remaining Medusozoa (*43*) and they lack all characters considered here.

We used a common assembly and filtering pipeline for public and newly-assembled transcriptomes and a separate pipeline to infer protein-coding genes from genomes, described below. A full list of reads and samples used in our dataset is in Data S1. The only two exceptions to these pipelines were from two recent publications from our group: a high-quality assembly for *Sarsia tubulosa* (*44*) and a recently-assembled genome of *Bougainvillia muscus* (*45*).

We lightly trimmed raw reads with Trimmomatic v0.39 (*46*) (TruSeq3-PE.fa:2:30:10 SLIDINGWINDOW:4:5 LEADING:5 TRAILING:5 MINLEN:25), performed kmer correction with RCorrector v1.0.6 (*47*) and removed ribosomal RNA with SortMeRNA v4.3 (*48*). We assembled transcriptomes using Trinity v2.15.1 (*49*) with default parameters and reduced redundancy by clustering transcripts at 99% identity using CD-HIT-EST v4.7 (*50*) (-G 0 -c 0.99 -aS 1.00 -aL 0.005). We predicted peptides using TransDecoder v5.7.1 (`https://github.com/TransDecoder/TransDecoder`), retaining peptides with hits to UniRef90 (downloaded March 8, 2024) searched by DIAMOND BLASTP v2.1.8 (*51*) in "sensitive" mode with evalue = $1 \times 10^{-5}$ or hits to PFAM (downloaded March 8, 2024) as searched by hmmscan (-E 1e-10; `http://hmmer.org`). Finally, we used CD-HIT to cluster 100% overlapping peptides (-c 1.0) and assessed transcriptome completeness with BUSCO v5.6.1 against the Metazoa odb10 database, downloaded January 11, 2024 (*52*). Protein models were annotated with EggNOG-mapper v2.1.12 (*53*).

For species with high-quality genome assemblies, we used RepeatMasker (`http://www.repeatmasker.org`) and Braker3 (*54*) to annotate protein-coding sequences. All accessions, including protein models and RNA-seq reads used to infer Braker3 gene models, are in Data S1.

Coding sequences were then clustered with CD-HIT as above.

We used an iterative approach to infer orthologs while minimizing potential errors due to contamination, homology inference, and sequencing artifacts. First, we used OrthoFinder v2.5.5 (*55*) with the settings "-y -M msa" to obtain Orthogroups (OGs) present in at least 30/98 species. For each OG, we used PREQUAL (*56*) (with settings '-filterthresh 0.9 -pptype closest 20') to mask non-homologous characters and re-aligned sequences using MAFFT-LINSI (for OGs with $\leq$ 1000 sequences) or MAFFT -auto (for OGs with > 1000 sequences). We used TrimAl (*57*) to trim sites with < 10% occupancy, and re-inferred gene trees using either 1) IQ-Tree v3.0.1 (*58*) with settings "-fast -mset LG+G -m TEST -alrt 1000" (for OGs with $\leq$ 1000 sequences); or 2) FastTree with settings "-LG -gamma -pseudo" (for OGs with > 1000 sequences). We then used TreeShrink v1.3.9 (*59*) with alpha=0.10 in 'all-genes' mode to remove potential paralogs, contaminants, or other erroneous sequences on long branches. We re-aligned the cleaned OGs using MAFFT-LINSI, trimmed sites with < 20% occupancy and sequences with < 50% overlap using TrimAl (-gt 0.2 -resoverlap 0.5 -seqoverlap 50), and dropped sequences with gaps in $\geq$ 80% of sites. We then re-inferred trees using IQ-Tree -fast -m Q.pfam+F+R10 and re-ran TreeShrink to remove any remaining long-branch sequences. We inferred gene trees one last time using IQ-Tree with Q.pfam+F+R10 (with the '-fast' flag for genes with $\geq$ 500 sequences).

To extract single-copy orthologs, we then collapsed nodes with less than 80% support into a polytomy and masked monophyletic tips from the same taxon by selecting the tip with the most non-ambiguous characters in the trimmed alignment. Finally, we used PhyloPyPruner `https://pypi.org/project/phylopypruner/` to infer 1-to-1 orthologs with the Maximum Inclusion algorithm (–min-taxa 30 –mask None –prune MI). Single-copy orthologs were again filtered with PREQUAL, aligned with MAFFT-LINSI and trimmed with TrimAl (-gt 0.2 -resoverlap 0.5 -seqoverlap 50), and we built final single-copy gene trees using IQ-Tree with the best-fitting model (-nt AUTO -B 1000 -bnni -alrt 1000 -msub nuclear). After trimming, we obtained 1771 orthologs present in at least 30 species. Symmetry testing in IQ-Tree revealed that 29/1771 loci (1.6%) violated assumptions of stationarity or homogeneity ($p < 0.05$), and we proceeded to analyze the remaining 1742 loci.

We built initial maximum-likelihood (ML) gene trees withIQ-Tree with best-fit models selected by ModelFinder, and inferred initial species trees using ASTRAL (*60*) and IQ-Tree; the IQ-Tree analysis was partitioned by gene. These approaches recovered the same topology (Fig. S10) and resolved all order-level relationships with maximal posterior support. However, some shallow relationships were unresolved because taxa contained little overlap in our data matrix (Table S1). To fill in family-level relationships, we also used ASTRAL-PRO3 (*60*) on the set of multi-copy gene families (OGs) containing less than 500 sequences (n=9194). ASTRAL-PRO successfully resolved the shallower relationships, recovered all genera as monophyletic (Fig. S11), and differed from the other trees only with respect to the placement of *Eudendrium carneum* with minimal support (55.1% posterior probability). We conducted subsequent analyses using the consensus topology and the placement of *E. carneum* from the ASTRAL and IQ-Tree phylogenies because of its high support (98.3% and 100%, respectively) and consistency with prior studies (*61, 62*).

To perform character mapping, we used the well-resolved phylogenomic topology as a backbone for a 621-taxon phylogeny (563 medusozoans and 58 anthozoans) derived from DNA barcoding sequences (*14*). To construct this phylogeny, we first identified gene sequences in our assemblies matching the 5 markers used by (*14*) (12S, 16S, 18S, 28S, and COI genes) and added them to the data matrix, using multiple rounds of constructing gene trees to identify and remove misaligned or misidentified sequences. We only analyzed species that were present in at least 2/5 gene alignments. We then used this DNA data matrix in a partitioned analysis together with 100 amino acid alignments in IQ-TREE. The amino acid alignments were the 100 best loci selected by kinda-date in the next section ('Divergence dating and fossil calibrations'). This mixed DNA+AA analysis was constrained to match the consensus topology from our species tree analyses above.

This analysis revealed that one of our new transcriptomes, originally identified as *Halecium sp.*, actually groups with *Nemalecium sp.* and is very likely *N. lighti*. Since we also had a transcriptome for *N. lighti*, we excluded this "*Halecium sp.*" sample from further analyses. We additionally dropped a sequence that was mis-labelled as "*Antennella secundari*" instead of the correct *Antennella secundaria*.

##### Divergence dating and fossil calibrations

We selected sequences for divergence dating using kinda-date (*63*), which sorts loci according to a weighted metric based on root-to-tip variance (clock-likeness), average branch length (information content), and Robinson-Foulds distance to the species tree (topological conflict). We constructed a

50-gene dataset based on these KINDA-DATE values (Data S2). In order to ensure sufficient coverage of the data-poor taxon *E. carneum*, we selected the top 5 genes containing *E. carneum* and the top 45 other genes overall. We constructed a 100-gene dataset in the same way (requiring at least 5 *E. carneum* genes), which was used to construct the 621-taxon phylogeny in the previous section. We then performed relaxed molecular clock analyses using MCMCTREE (*64*), including a series of sensitivity analyses considering variation in rate priors, gene sampling, and taxon sampling (see Supplementary Text).

We chose between independent- and correlated-rates models using model selection based on marginal likelihood calculations, a procedure recently developed for approximate likelihood methods using branch length transformations within MCMCTREE (*65*). We fixed the root age at 1 without fossil constraints for the purpose of marginal likelihood estimation. We analyzed the 68-taxon 50-gene dataset using 32 beta samples and ran chains sampling every 50 iterations for 800,000 samples, with a burn-in of 100,000 iterations. The independent-rates model ('usedata = 2') was strongly preferred (PR=0.997 vs. 0.003), so we used this model for all analyses.

To run MCMCTREE, we used the coalescent species topology and estimated Hessian matrices and branch lengths under the LG substitution matrix and a five-category gamma model of rate variation. We ran each MCMCTREE analysis with two chains, each sampling every 500 iterations for 20,000 samples following a burn-in of 1,000,000 iterations, and also performed MCMC sampling from the prior using the same settings.

All fossil constraints are 2.5% soft minima unless otherwise noted. Lower bounds as implemented in MCMCTREE take the form of truncated Cauchy distributions. For constraints with minimum ages < 300 Mya, we used the default scale parameter of 1 (providing a broad distribution), while we used a scale parameter of 0.25, which places more of the prior density close to the minimum bound, for ages > 300 Mya. The root was constrained with a skew-normal distribution with a hard minimum and a 5% soft maximum such that the 0.95 quantile was 609 Mya; see Data S3 for details on calibrations.

Finally, we transferred divergence times from MCMCTREE to the 621-taxon phylogeny using CONGRUIFY (*66*) and TREEPL (*67*). We did this for the mean posterior MCMCTREE as well as for 500 trees sampled from the posterior to account for divergence time uncertainty. In TREEPL, we set 'thorough = true' and 'numsites = 125892' (the number of sites in our DNA+AA alignments) and followed the procedure of (*68*). We first used 'prime' 3 times to select the best optimization parameters (opt = 2, optad = 2, optcvad = 2); then used 'randomcv' 3 times to select the best smoothing parameter ($1 \times 10^{-11}$); and finally ran TreePL with these parameters to produce the time-calibrated tree.

**Ancestral state reconstructions**

We conducted literature searches for eye presence/absence, medusa/gonophore morphology, and colony architecture (Data S4). For eyes, we followed (*14*) in defining eyes minimally as a "region made of photoreceptor cells adjacent to pigment cells." This definition includes simple ocelli up to the complex lensed eyes of Cubozoa. Although Miranda & Collins (*69*) discuss potential eyes in some staurozoans due to the presence of rhopaliar pigment spots, we scored these species as uncertain ("?") because there is no evidence that these regions contain photoreceptors.

Hydrozoans exhibit a continuum of reduced medusa morphologies. The gonophore (an asexual bud that becomes the medusa) may be reduced or completely lost, and if present may be liberated from or attached to the colony. We used the following gonophore encoding scheme based on Cartwright & Nawrocki (*16*) and Miglietta & Cunningham (*15*): 1) motile, autonomous feeding medusae, 2) non-feeding medusoids with radial canals and/or tentacle bulbs, 3) gonophores lacking radial canals and tentacle bulbs (sporosacs), and 4) gonophores absent. Under these definitions, medusoids and sporosacs may still detach briefly from the colony but are non-feeding, and Category 1 includes the benthic medusae of *Staurocladia spp.* and Staurozoa. We also tested binary models of medusa presence, where categories 2, 3, and 4 above were all encoded as 'absent.' To provide realistic models of character evolution, we disallowed re-gains of medusae from any other state (*15*).

For colony architecture, species were scored as: 1) lacking a polyp stage, 2) solitary, 3) encrusting colonies, 4) upright colonies, and 5) pelagic colonies. We did not allow polyps to be re-gained once lost. Additionally, based on initial analyses, we found that upright and pelagic colonies were never lost and therefore also disallowed losses of those states.

We inferred ancestral state reconstructions (ASRs) using corHMM v2.10 (*70*) with the 'maddfitz' root prior and 10 random restarts. We tested a variety of models with up to 4 hidden rate categories and reported the best-fitting models in the main text (Data S4). We identified conservative minimum numbers of character transitions by manually inspecting node marginal likelihoods and uncertainty (bootstrap support) in the species tree. Specifically, we considered nodes with bootstrap support < 80% as unsupported; if an ASR inferred multiple origins separated by such an unresolved node, we conservatively consider this a single origin. We inferred ASRs on a phylogeny without Anthozoa because anthozoan colonies are not homologous to medusozoan colonies (*71*) and our encoding scheme does not apply to them. To estimate ages of character transitions, we examined the posterior distributions of relevant node ages from 500 randomly sampled posterior trees.

We tested for phylogenetic correlations between characters using the test proposed by Boyko & Beaulieu (*72*), which involves comparing fits of independent and correlated models with both 1-rate Markov models and 2-rate hidden state models (HMMs). We considered an evidence ratio (ER) > 2.7 as supporting one model over another, and found significant correlations between eyes and medusae, and between colony type and medusae. We report the results of the joint eye-medusa model in the main text. We found that there was no difference in the number of inferred colony type transitions regardless of modelling it jointly with medusae (see Supplementary Text).

**Quantifying adaptive convergence**

We used CSUBST (*12*) to infer convergence events between each pair of species (n=86 medusozoans, n=3570 species pairs) using the set of 9194 OGs used for the ASTRAL-PRO analysis. Gene trees were midpoint rooted and codon sequences were obtained using PAL2NAL v14.1 (*73*). After running CSUBST in 'analyze' mode on each gene tree and codon alignment with setting '–max_arity 2,' we extracted all branch pairs with at least 3 non-synonymous convergent substitutions ($OCNany2spe \geq 3$) and $omegaCany2spe \geq 3$, as recommended by the CSUBST authors, and considered these branch pairs "convergent." To subset data for the eye, medusa loss, and coloniality data, we extracted only the comparisons between phenotypically convergent lineages identified by ASR. For medusa-loss species, we further excluded medusoids and only considered comparisons between species with either sporosacs or no gonophores. For colonial species, we considered comparisons between species with upright colonies.

See the Supplementary Text for analyses and discussion of technical considerations for how we scored convergence between species pairs.

**Regression analyses**

For each pair of species, we calculated the proportion of OGs with at least one convergence event between them and performed phylogenetic regression analyses to test for a relationship with divergence time. We accounted for phylogenetic structure using the recently-developed approach of Anderson et al. (*13*) to calculate the phylogenetic covariance matrix for species pairs, with the 'taxapair.vcv' function of their PHYLOPAIRS R package. We then used this covariance matrix as input for phylogenetic generalized least-squares (PGLS) analyses with the 'gls' function of the NLME R package. Convergence values were right-skewed, so we used a Yeo-Johnson transformation (*74*) on convergence values for all subsequent analyses; though untransformed values gave similar results (Fig.S12). Results were similar if we calculated convergence using a more stringent statistical cutoff ($\omega_c \geq 5$) or considered the total number of convergence events per OG rather than scoring each OG as either convergent or not (Fig. S13).

In addition to divergence time, we considered protein distance and transcriptome size as covariates. As our measure of protein distance, we calculated the median pairwise protein distance between each pair of species using the 'dist.alignment' function of the SEQINR R package (with the matrix='similarity' argument); this calculates the Fitch similarity matrix (*75*). These values were normalized within each OG using a modified Z-score (which uses the median rather than the mean) based on the median protein distance across all species within the OG. This accounts for baseline differences in evolutionary rate among OGs. These normalized protein distances were strongly correlated with divergence time ($r = 0.90$, Fig. S18). For transcriptome size, we simply calculated the mean transcriptome size of the two species (which was not correlated with divergence time; $r = -0.077$, Fig. S18). We then tested all possible models containing divergence time, protein distance, and transcriptome size, and found that the full model with all interaction terms was by far the best fit (Table S5). We analyzed standardized regression coefficients using the 'plot_model' function of the SJPLOT R package.

To test for differences between eye, medusa loss, and coloniality data subsets, we included these grouping factors as covariates in the model and tested for group differences and interaction terms. Because these categories were non-exclusive (14 species belonged to both the "medusa loss" and "upright colony" groups), we coded each group as a binary variable and ran a regression analysis as follows: [$cts \sim Div\_time * (is\_Eye + is\_Med + is\_Col)$]. This model showed that the group intercepts and interaction terms were all non-significant ($p > 0.1$), and standardized regression coefficients revealed no groupwise differences (Fig. 2C).

To test whether certain taxa systematically differed in levels of convergence, we averaged convergence values within each species and asked whether these values had phylogenetic signal (using the 'phylosig' function of the PHYTOOLS R package) or differed among clades. With raw convergence values, species with larger transcriptome sizes had more convergence, resulting in higher convergence values on average within Hydrozoa (Fig. S8). These values had a marginally significant phylogenetic signal ($p = 0.037$, Blomberg's k, 10,000 simulations). However, analyzing residuals of the full regression model removed this pattern and resulted in no phylogenetic signal ($p = 0.78$, Fig. S8).

**Null model of convergence**

Inspired by the Zero-Force Evolutionary Law (ZFEL) (*19*), we used two-state Markov chains to model the expected decline of convergence over time in the absence of systematic forces or constraints. Our null hypothesis is that all gene families are equally likely to be selected; specifically, 1) all convergence is the result of random (uncorrelated) overlaps between selected genes in different species, and 2) there are no constraints on convergence across genes. Many parameterizations of this null model are possible, but we can estimate an upper bound by assuming that the null cannot exceed observed convergence levels.

We defined a ZFEL-like model of convergence as follows. Consider a genetic locus (here, a gene family) as a binary variable with 1 indicating selection. A locus is convergent between two species if its value is 1 in both. At each time step, a locus can transition from 0 to 1 with probability $P_{gain}$ or from 1 to 0 with probability $P_{loss}$. This is a two-state Markov chain with steady-state frequency of [ $S = P_{gain}/(P_{loss} + P_{gain})$] (*76*). Given two vectors of $n$ loci, the expected proportion of overlap due to random chance is simply the product of the proportions of ones in each vector, which at equilibrium equals $S^2$. Given $P_{loss}$, $P_{gain}$, and a starting proportion of ones $S_0$, we can use random walks to simulate convergence over time, which will eventually reach the steady state. For a decline over time to occur, $P_{loss} > P_{gain}$ and $S_0 > S$. Biologically, the null hypothesis of our ZFEL-like model is that all loci are equally likely to be selected on average; specifically, 1) there are no constraints on convergence across loci, and 2) selection does not act to systematically drive convergence (all convergence is the result of random overlaps between loci).

$P_{gain}$ and $P_{loss}$ are unknown because we do not know the proportion of selected loci in each species. Nonetheless, we can estimate a conservative upper bound on convergence under the null expectation as follows, assuming a single $P_{gain}$, $P_{loss}$, and $S_0$ for all species. First, we make the key assumption that null expectations cannot exceed observed convergence levels, because that would require invoking selection against convergence (e.g., character displacement) even among species with no spatiotemporal overlap. Observed convergence levels can then be considered a theoretical upper bound on the null. We estimate $S_0$ as the square root of the fitted divergence time regression at time 0 ($S_0$ = 0.3878). This is the number of "selected" loci that must exist on average to explain the observed level of convergence, assuming all overlap is simply due to chance. We then assume that S is the square root of the mean value at the oldest time point in our dataset, 681.94 Mya ($S$ = 0.201).

For a given value of $S$, there is a defined relationship between $P_{gain}$ and $P_{loss}$ [$P_{gain} = (S * P_{loss})/(1 - S)$] (*76*), but these rates are not uniquely estimable. Therefore, we used simulations to estimate the smallest possible $P_{loss}$ consistent with observed levels of convergence given our estimates of $S_0$ and $S$. Specifically, we performed 500 simulations on vectors of length n=4501 (the mean number of shared OGs per species pair) over a range of $P_{loss}$ values (Fig. S14) and selected the largest value for which $S$ fell within the 95% confidence interval of the simulated data at the 681.94 My time point. This value was $P_{loss} = 0.00310$ (Fig. S14), which requires $P_{gain} = 7.789 \times 10^{-4}$. We then used these parameters to simulate expected amounts of convergence between each species pair based on their divergence time and number of shared loci.

To represent the fact that our chosen $P_{loss}$ is the minimum of a range of possible values, we also simulated data under three larger values of $P_{loss}$ (0.0035, 0.005, and 0.01) that are visualized in Fig. 2B. For each value of $P_{loss}$, we binned data into 50-Myr intervals and compared real and simulated convergence using paired t tests weighted by the inverse variance of the simulated data. P-values were Bonferroni-corrected within each $P_{loss}$.

**C-scores and gene ontology analyses**

C-scores were calculated for pairs of species pairs (quartets). Specifically, we calculated the overlap of convergent genes as the ratio of OGs that independently experienced convergence in both species pairs to the set of all OGs found in both species pairs, and calculated C-scores from the hypergeometric distribution (*28*). To ensure that we only considered independent convergence events, we only counted convergence events that occurred along different branches of a gene tree. This procedure is computationally intensive because we have to map substitutions onto gene trees. Rather than analyze all 4,755,188 possible non-overlapping quartets, we drew 50 random samples of 100 species pairs each, resulting in 247,500 total quartets (236,070 after excluding species overlaps). We then used the covariance between species pairs in the lineage-pair phylogenetic covariance matrix as a measure of phylogenetic distance and performed a linear regression between C-scores and covariance using R's 'lm' function.

We performed GO enrichment analyses of convergent genes (in the 'Biological Process' category) for each species pair (n=3570), where the denominator ("gene universe") consisted of all OGs shared between those species and passing the $OCN \geq 3$ cutoff (see Supplementary Text). We used TopGO (*77*) with Fisher's exact test and 'weight01' algorithm and considered GO terms significant at a level of $p < 0.001$. We calculated the semantic similarity between all combinations of the 2082 unique GO terms using REVIGO (*78*). We then compared lists of GO terms for all 4,755,188 species quartets using the 'Best Match Average' method (*79*), which is the mean of the maximum similarity between each GO term and members of the other list. We regressed these values against phylogenetic covariance using 'lm', as above.

We also performed GO enrichment at the OG level across all species using the 'KS test' functionality of TopGO; we used the per-gene convergence rate (total number of convergence events divided by the number of genes) as input. We performed separate tests in each direction to detect enriched terms among the most- and least-convergent OGs. Fig. S9 shows the most significant GO terms for this analysis, and Data S5 lists all significant terms ($p < 0.01$).

Finally, we asked whether genes with known roles in phototransduction and cnidarian eye development experienced higher rates of convergence among eye-bearing species. We compiled two sets of genes: 1) homologs of differentially-expressed genes (DEGs) upregulated in the eye-bearing tissues of *A. aurita*, *T. cystophora*, and *S. tubulosa* (*44*), and 2) gene families belonging to a previously-defined list of light-interacting genes (*80*). We added a handful of genes known to play roles related to light perception in cnidarians (Pax transcription factors and a hydrozoan eye opsin; (*44*)) and modified the set of circadian-related genes to reflect current knowledge of clock genes in Medusozoa (*81*). In total, we identified 798 eye-associated OGs belonging to these gene sets.

**KEGG gene set enrichment analysis**

We used KEGG pathway annotations to classify certain gene sets as either "environment-facing" or "housekeeping." This allows us to test whether these *a priori*-defined gene sets differ in their levels of convergence, orthogonal to the above exploratory analyses based on GO terms. Specifically, we retrieved OGs annotated with KEGG terms we considered to fall unambiguously into these two categories (Table S4). We then ranked OGs according to their per-gene convergence rate as above and used the 'GSEA' function of the clusterProfiler R package (with pAdjustMethod='BH', maxGSSize = 10000, and sampleSize=10000) to test for enrichment of these two gene sets. For visualization purposes, we also calculated enrichment of the individual KEGG categories (Fig. 3D).

**Physicochemical classes of convergent substitutions**

Although CSUBST seeks to identify signatures of positive selection, $\omega_c$ could also be elevated in situations when site substitution profiles are constrained and multiple amino acids at that site are functionally equivalent (*12*). In that case, we expect substitutions to produce amino acids with similar physiochemical properties. If convergence is instead driven by positive selection, we expect substitutions to more frequently alter physicochemical properties. To test these possibilities in our data, we analyzed all double substitutions—where both genes experienced a change in amino acid state—on branch pairs identified as convergent between eye-bearing species. Double substitutions may or may not be convergent.

We first grouped amino acids based on physicochemical similarity using four alternative classification schemes (GS1-4 from (*82*)). We then categorized substitutions as either physicochemically conservative (meaning one or both substitutions occur within the same class) or physicochemically convergent (meaning both substitutions change physicochemical class, resulting in the same derived state). We only analyzed substitutions where both derived amino acids belonged to the same physicochemical class, in order to make a fair comparison with convergent substitutions. For each category, we tested whether the proportions of conservative substitutions differed between convergent and non-convergent double substitutions using a chi-squared test. We repeated this analysis for the "false positive" data simulated under zero convergence (Fig. S7).

#### Supplementary Text

##### Divergence dating sensitivity analyses

We assessed the effects of prior parameters by systematically varying the priors for mean substitution rate (rgene-gamma), rate heterogeneity (sigma2-gamma), and the sampling parameter of the birth-death process (BDparas) on the 28-taxon dataset. Initial parameters were "rgene-gamma = 2 10", corresponding a mean substitution rate of 0.2 substitutions per 100 My, "sigma2_gamma = 2 10" corresponding to high rate heterogeneity among lineages, and "BDparas = 1 1 0.1." Posterior results were largely insensitive to variation in the birth-death and Dirichlet (rgene-gamma) priors (Fig. S4). The parameter with the largest impact was the rate drift parameter sigma2, which controls rate variation among lineages, though its effect was also small. Reducing the mean of the rate drift prior by an order of magnitude (sigma2_gamma = 2 100) caused older posterior ages at crown Cnidaria (679 vs. 648 Mya) and Medusozoa (675 vs. 643 Mya). These times may be biased upwards because failure to accommodate among-lineage rate heterogeneity negatively impacts divergence time estimates. In contrast, increasing this parameter by an order of magnitude (sigma2_gamma = 2 1) resulted in only slightly younger times at old nodes (640 Mya for Cnidaria, 634 Mya for Medusozoa). This suggests that our original choice of prior adequately accommodated rate heterogeneity and did not strongly bias our results.

There was a moderate effect of taxon sampling on posterior age estimates, with higher taxon sampling resulting in older divergence times (Fig. S5). Increasing taxon sampling should generally improve divergence time estimates, both because poor taxon sampling can underestimate node ages (*83*) and because better taxon sampling enables the use of more fossil information. Therefore, we favour the largest 68-taxon dataset. Note that we still recover a Cryogenian origin of Medusozoa even with 28 taxa (643 Mya).

We also analyzed the second-best set of 50 genes identified by KINDA-DATE, and found that gene sampling had a much smaller effect on divergence times than taxon sampling. Overall, posterior distributions had extensive overlap across all sampling schemes (Fig. S5).

##### Correlations between characters

Accounting for correlations between phylogenetic characters is important to accurately account for transition rates. We investigated this in our data because we expected strong correlations among our characters (e.g., eyes are only found on medusae). We found a significant correlation between eyes and medusae ($ER = 1.5 \times 10^{12}$) and between colony type and medusae ($ER = 410$), but not between eyes and colony type ($ER = 2.3 \times 10^{-6}$; Table S6). Since medusa presence/absence was strongly correlated with both other characters, we conducted joint reconstructions of medusae and eyes, and medusae and colonies, in addition to individual reconstructions of each character. For eyes, we found that the number of inferred origins was sensitive to the decision to model jointly with medusa state (Fig. S16; 9 inferred origins in the joint model, 12 in the independent model). Thus, failure to account for the strong trait dependence between eyes and medusae can mislead estimates of the rates of eye gain and loss. We therefore report the results of the joint model. On the other hand, the number of inferred colony type transitions was identical regardless of whether or not it was modelled jointly with the medusa stage.

##### Calculating convergence between species pairs

Convergence events can occur along terminal or internal branches. Thus, we need to propagate convergence events occurring on internal branches in order to quantify convergence between pairs of extant species. Consider a hypothetical convergence event occurring between two internal branches as in Fig. S2, each with two descendant species. This event is supported by 3 convergent amino acid substitutions, and there are 4 pairwise species comparisons affected by this convergence event. Fig. S2B shows how each pairwise comparison is scored according to different possible scoring schemes. In the "simple" scheme (used in the main analyses), we consider all comparisons as convergent because a convergence event occurred between their ancestors, regardless of whether the convergent residues are conserved in extant sequences. Other scoring schemes require a minimum number of the convergent ancestral residues to be conserved. In practice, checking the extant sequences requires running CSUBST in 'sites' mode, which is computationally demanding. We used the eye- and medusa-loss data subsets to compare these scoring schemes and found that they made no difference to the overall results (Fig. S2C) and affected a negligible number of branches. Therefore, we used the "simple" scoring scheme for the main analyses.

**Phenotype-focused vs. phenotype-blind convergence**

When scoring convergence between species sharing a convergent trait or ecological regime, most studies only consider comparisons between branches after the inferred origin of that trait (often referred to as "foreground" branches). For instance, if we consider the comparison between two foreground species in Fig. S3, we would compare only the blue branches. This makes sense if the goal is to identify genes that may be associated with the trait(s) of interest, but excludes convergence that may have taken place between the ancestors of the two species for other reasons.

In our case, we quantified convergence among all species pairs irrespective of any phenotype, meaning that there are no foreground or background branches. When subsetting the data to focus on phenotypically convergent lineages, we quantified convergence using both the above phenotype-focused approach (Fig. S9A, blue dashed lines) and the phenotype-blind approach (blue and grey dashed lines). In the main text, we used the phenotype-blind approach for all analyses in order to facilitate statistical comparisons between these data subsets and the full dataset. However, there was no statistical difference between phenotype-blind and phenotype-focused convergence in any of the three phenotype data subsets (eyes, medusa loss, upright colonies; Fig. S3).

**Requiring a minimum number of non-synonymous substitutions biases convergence measures if not accounted for**

Like traditional dN/dS scans, $\omega_c$ cannot reliably detect selection if too few substitutions have occurred (e.g., (*84*)), so the authors of CSUBST recommend requiring a minimum number of non-synonymous convergent substitutions in order to identify a branch pair as convergent ($OCN \geq 3$) (*12*). This means that detection of convergent branch pairs is biased against similar sequences and, since convergent substitutions are rare, this could potentially confound a relationship with divergence time. We found that this was indeed the case. In a naive analysis, we calculated the proportion of convergent OGs using the total number of OGs shared between species pairs, which resulted in a positive relationship between convergence and divergence time (Fig. S1A). This occurred because the proportion of OGs passing the OCN cutoff was strongly correlated with divergence time ($r = 0.75$, Fig. S1B). To account for this bias, we only included an OG in a species pair's denominator if it contained at least one pair of proteins separated by at least 3 convergent nonsynonymous substitutions (OCN) along at least one branch. In other words, we only considered OGs where it was theoretically possible to detect convergence based on the minimum number of substitutions; doing this resulted in the expected negative relationship between convergence and divergence time found in the main text.

##### Simulations to estimate false positive rate

To estimate an empirical false positive rate and assess potential biases in our gene alignments, we simulated codon evolution along gene trees under scenarios of zero convergence, with parameters and alignment lengths derived from the real alignments ('csubst simulate'). There were 24,933 false positives in the simulated data for a false positive rate of $5.31 \times 10^{-5}$. Assuming the same number of false positives in the real data, this corresponds to a reasonable false discovery rate of 0.059. Simulated counts had no relationship with divergence time (PGLS, $p = 0.4$; Fig. S6A). These data provide further evidence that observed patterns of convergence are driven by selection rather than neutral processes or biases present in the gene alignments. False positives in alignments simulated without convergence may reflect inaccuracies in the underlying codon substitution model and gene tree, or errors in the ancestral state reconstructions. In the full regression model of simulated false positive convergence, there was no effect of divergence time, while transcriptome size still exerted a slight positive effect (Fig. S6B). Protein distance had a positive, not negative, effect on simulated false positives, presumably due to the increased likelihood of errors in more divergent sequences. The fact that real data showed the opposite trend, in line with theoretical expectations, indicates that CSUBST primarily identifies genuine convergent substitutions. Simulated values had a much lower number of sequences passing the minimum $OCN \geq 3$ cutoff (as expected based on the lower amount of non-synonymous convergent substitutions), resulting in simulated convergent proportions actually being generally higher than observed values. This makes no difference in terms of inference, and the much greater OCN values in the real data support a signal of elevated biological convergence not attributable to biases in the underlying gene alignments.

#### Null model(s) of convergence

As discussed in the main text and Methods, $P_{loss}$ and $P_{gain}$ are not uniquely identifiable and there are thus infinite possible ways of implementing a null model. Nonetheless, there are two main reasons why our implementation is ultra-conservative (darkest null line in Fig. 2B). First, we chose the steady-state frequency $S$ to match the end of our time series, but convergence is likely to continue declining on longer timescales. Second, our initial value $S_0$ is based on the unrealistic assumption that all overlaps, even among closely related taxa, are due to random chance rather than shared genetic constraints or selection. If this assumption is violated, then the true value of $S_0$ (the proportion of "selected" loci that are capable of convergence) will be lower, but never higher. Additionally, since we chose our value of $P_{loss}$ using the 95% confidence interval of simulated values (rather than the mean), we are effectively testing our data against the upper 95th percentile of realizations of the null model (explaining why the oldest time points in Fig. 2B fall below the null line). In effect, we have constructed a null model to approximate the observed data as closely as possible, and still clearly reject the null.

#### Convergence in eye-related genes

Eye-related genes had relatively higher rates of convergence than other gene families among all species, but not specifically among eye-bearing species (LME, $p = 0.2$; Fig. S15). Thus, known eye-related genes are not strongly enriched for convergence between species with eyes. Nonetheless, GO enrichment of OGs with convergence in multiple eye-bearing lineages recovered significant terms related to phototransduction and eye development, including "embryonic camera-type eye morphogenesis," "circadian behavior," and "cellular response to retinoic acid" (Fig. S15). Additionally, two notable transcription factors with known roles in cnidarian eye development experienced convergent selection across 4 out of 5 eye origins: Pax (OG0000843) and Six (OG0001052) (Table S2). This might suggest that substitutions in these regulatory genes are implicated in eye evolution. However, many species without eyes also had convergent substitutions in both Pax and Six (Table S2). Further inspection revealed that every OG with convergence in three or more eye-bearing lineages also experienced at least one convergence event between species without eyes. We conclude that, although focusing on convergence among multiple eye-bearing lineages might slightly enrich for trait-related genes, homologs of those genes also frequently experience convergence among species lacking that trait.

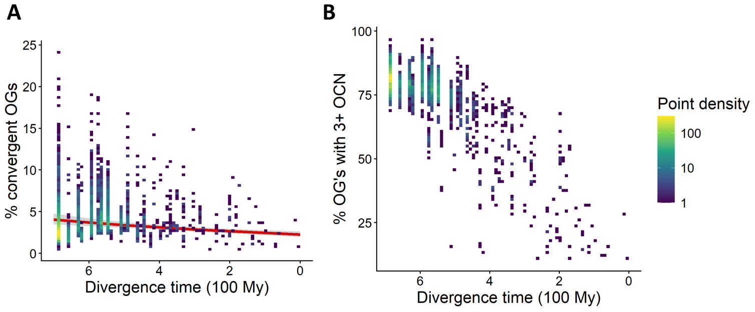


**Figure S1**: **Failure to account for minimum number of convergent substitutions biases convergence estimates.** A) Phylogenetic regression of convergence against divergence time including all OGs shared between species, without accounting for whether sufficient substitutions have occurred to detect convergence. B) The proportion of OGs with sufficient substitutions ($OCN \geq 3$) is strongly biased towards older species pairs.

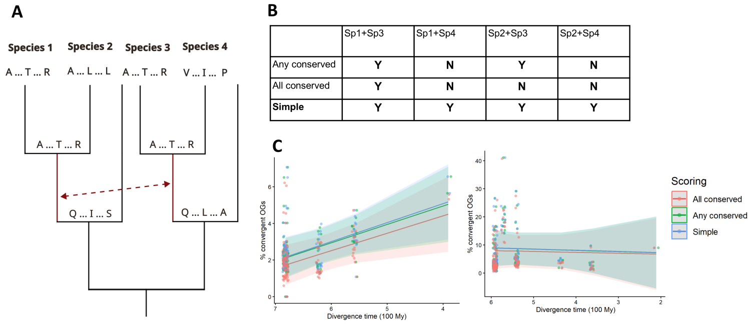


**Figure S2**: **Propagation of convergence events to terminal branches does not affect inferred patterns of convergence.** A) Schematic of hypothetical convergence event between two internal branches (red arrow). B) Table showing convergence between extant species pairs under alternative scoring schemes. 'Y' indicates scored as convergent, 'N' indicates scored as non-convergent. "Any conserved" requires at least one conserved extant residue; "All conserved" requires all residues to be conserved; and "Simple" has no such requirement. C) Phylogenetic regressions of convergence against divergence time under these scoring schemes, for eyes (left) and medusa loss (right). All slopes and intercepts were statistically indistinguishable ($p > 0.1$).

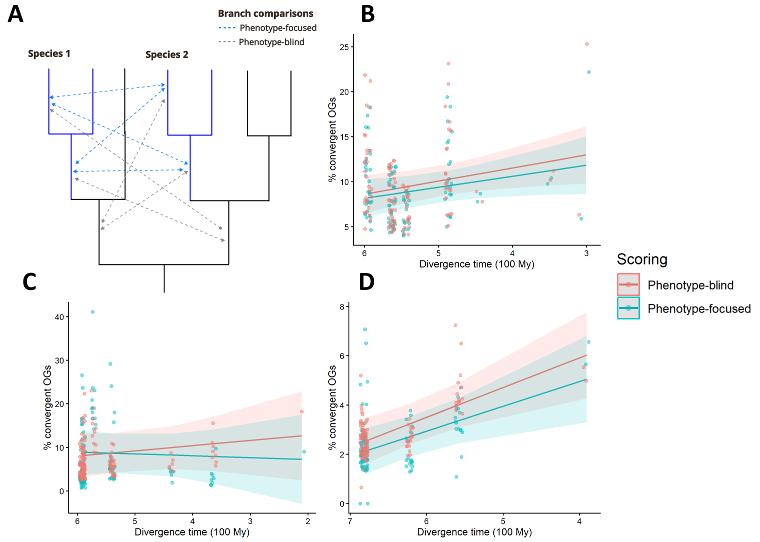


**Figure S3**: **Excluding branches based on ancestral phenotypes does not affect inferred patterns of convergence.** A) Schematic showing all possible branch comparisons between two species sharing a convergent trait (blue branches). Comparisons between foreground branches (after the trait is inferred to have evolved) correspond to the phenotype-focused approach, while the phenotype-blind approach includes comparisons among other branches. B-D) Phylogenetic regressions of convergence against divergence time for both approaches, for eyes (B), medusa loss (C), and upright colonies (D). All slopes and intercepts were statistically indistinguishable ($p > 0.1$).

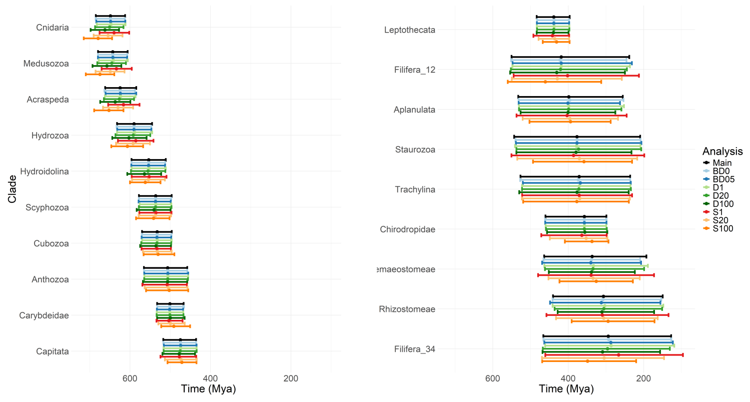


**Figure S4**: **Posterior clade ages are insensitive to variation in rate priors.** Analyses were performed with 28 species and 50 genes. Error bars represent 95% HPDs. BD: Birth-death prior, with tested values 0 and 0.5. D: Dirichlet (rate variation) prior with tested values 1, 20, and 100. S: Sigma_2 (among-lineage rate variation) prior with tested values 1, 20, and 100. "Main" is with parameters BD = 0.1, D = 10, and S = 10.

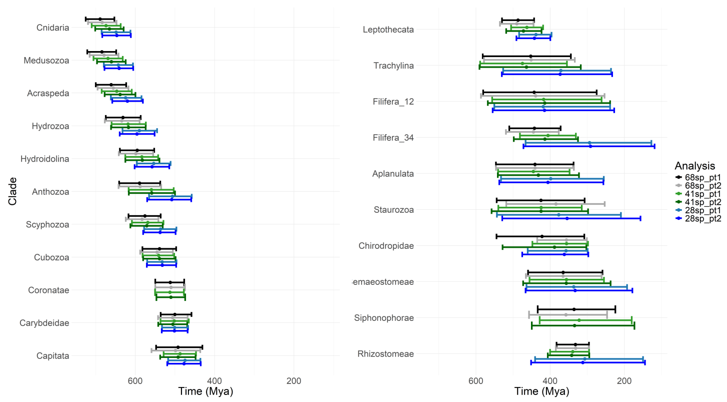


**Figure S5**: **Minimal effects of gene and taxon sampling on divergence time estimates.** Analyses were performed with 68, 41, or 28 species subsampled to capture major clade (Siphonophorae was not included in the 28sp analysis), and two alternative sets of 50 genes each. Error bars represent 95% HPDs. "_pt1" refers to the set of 50 genes used in the main analysis, while "_pt2" refers to the second-best 50 genes as selected by kinda-date (again requiring at least 5 genes representing *E. carneum*). All trees and result files can be found on the Github and Dryad.

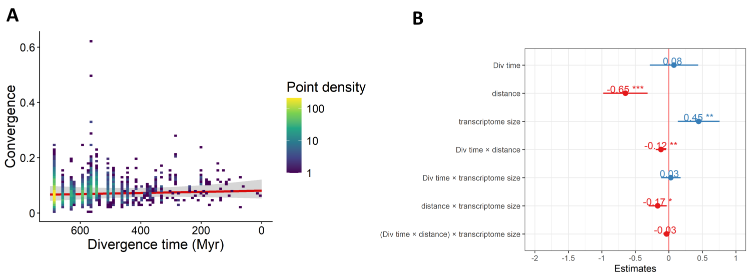


**Figure S6**: **Simulated false positives have no relationship with divergence time.** A) Phylogenetic regression of false-positive convergence values detected by CSUBST in data simulated without convergence (see Supplementary Text). B) Standardized regression coefficients of the regression with all covariates using the simulated data.

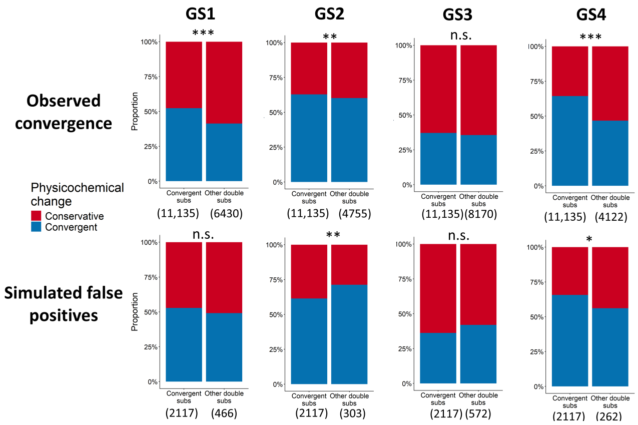


**Figure S7**: **Convergent amino acid substitutions identified by CSUBST are more likely to alter physicochemical properties than other substitutions.** We analyzed all branch pairs with convergence between eye-bearing species (top row) and identified all double substitutions (see Methods). We classified substitutions as physiochemically conservative or convergent based on four alternative encoding schemes (GS1-4). We performed the same analysis for simulated false positive convergence between eye-bearing species (bottom row). Numbers in parentheses indicate sample size; this value differs for other double substitutions because we excluded substitutions resulting in different physicochemical classes (see Methods). *: $p < 0.05$. **: $p < 0.01$. ***: $p < 0.001$ (Bonferroni-corrected p-values).

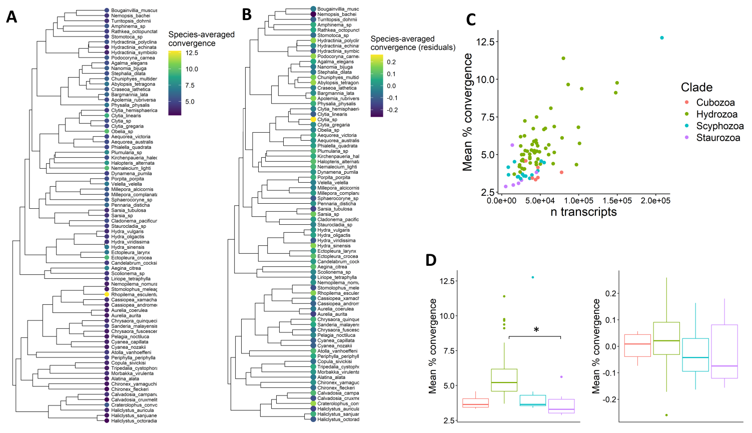


**Figure S8**: **Species with larger transcriptomes have higher levels of convergence, but there is no phylogenetic pattern after accounting for this.** A) Species tree showing species-average convergence values. B) Species showing species-average convergence values after accounting for transcriptome size (residuals of phylogenetic regression). C) Relationship between convergence and transcriptome size; note that hydrozoans tend to have larger. transcriptomes. D) Convergence values plotted for each clade, using raw convergence values from the tree in A (left) and residual values from the tree in B (right). The only significant difference was between Hydrozoa and Staurozoa in the left-hand plot.

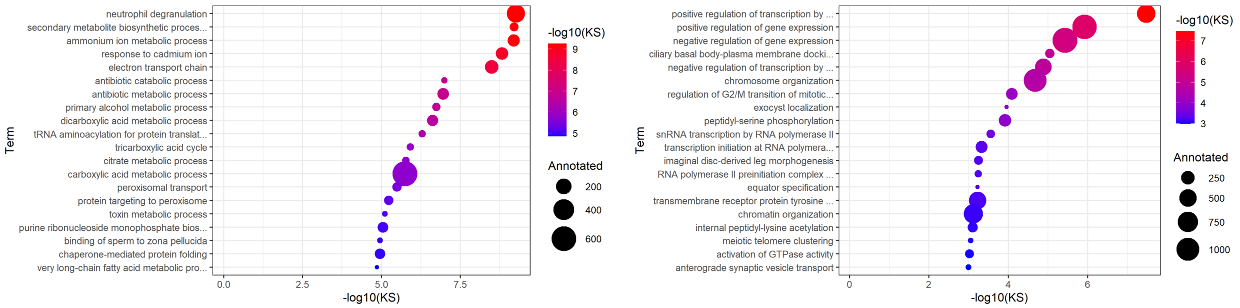


**Figure S9**: **Top GO terms enriched among most- (top) and least- (bottom) convergent OGs.** OGs were ranked according to their per-gene convergence rate (n convergence events divided by n genes) across all species. "KS" is the raw p-value from the Kolmogorov-Smirnov test implemented in TopGO. "Annotated" refers to the number of OGs annotated with a corresponding GO term. Full lists of GO terms are in Data S5.

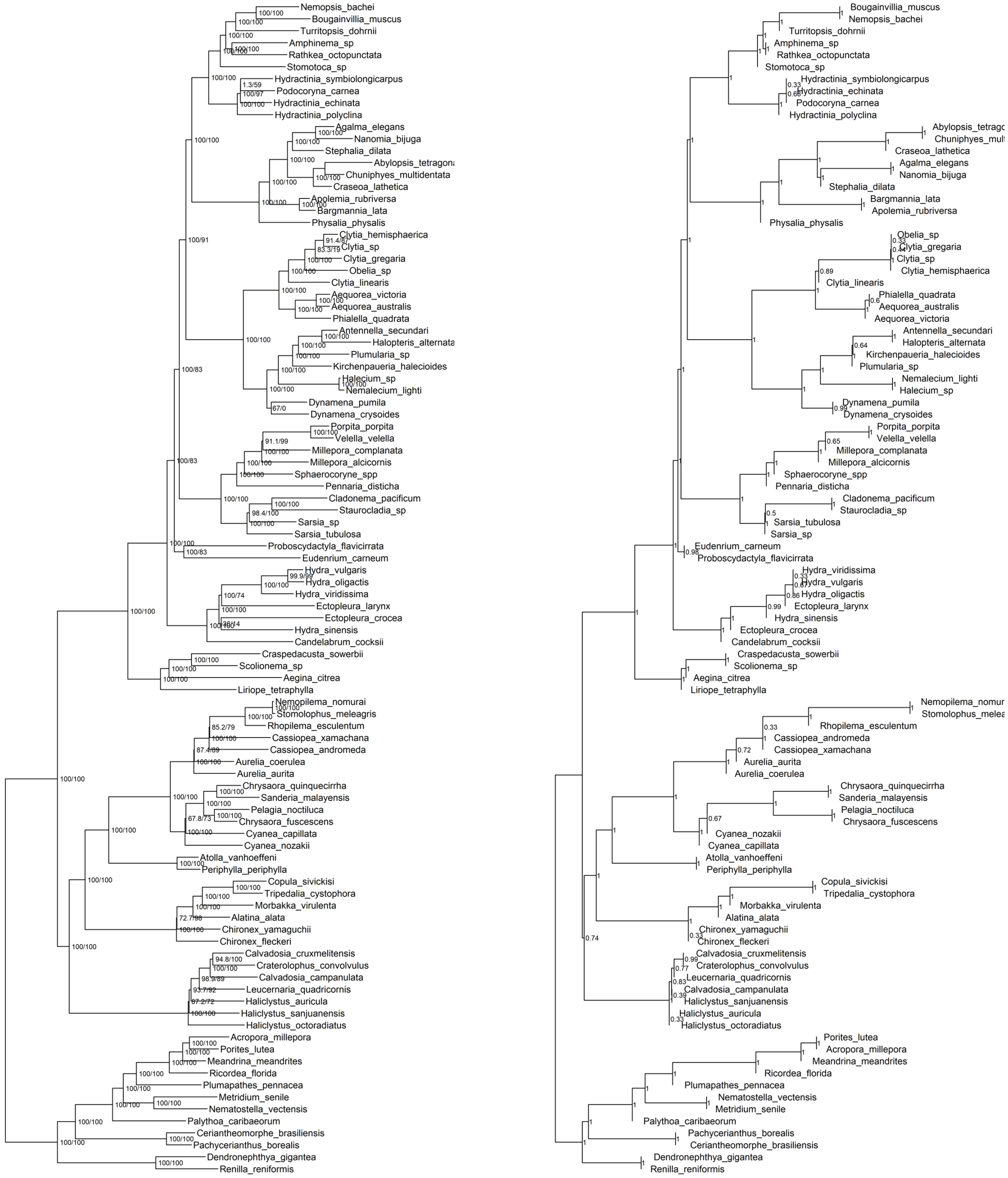


**Figure S10**: **Species trees inferred with IQ-TREE (left) and weighted ASTRAL (right) from 1742 single-copy orthologs.** Numbers indicate branch support values: ultrafast bootstraps and SH-aLRT for IQ-TREE (left to right, respectively) and quartet scores for ASTRAL. The topologies are identical except for some unresolved shallow relationships, which reflects the lack of overlap between these sequences in the data matrix (Table S1).

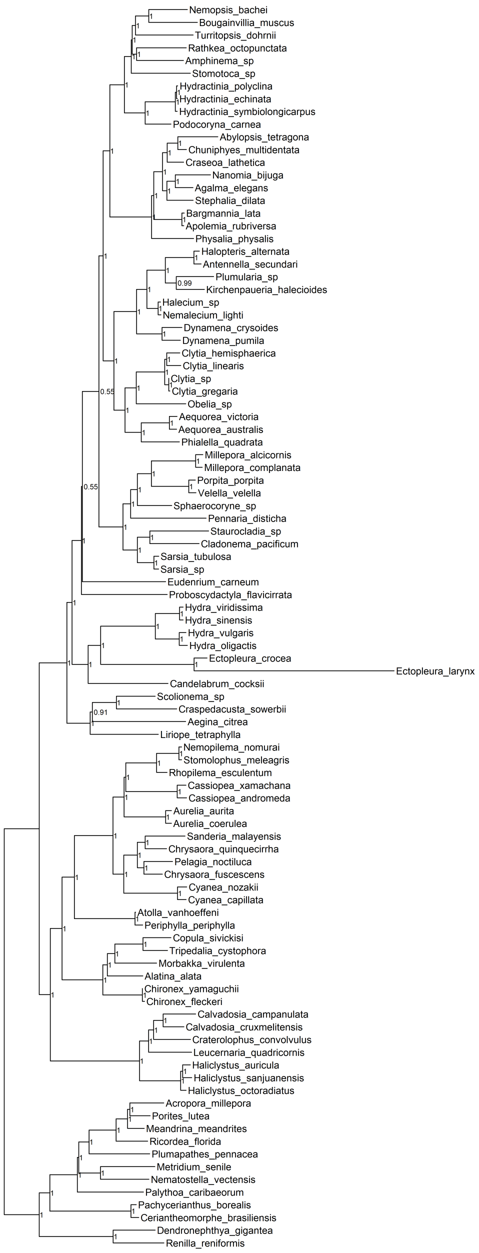


**Figure S11**: **Species tree inferred with ASTRAL-PRO from 9194 multi-copy orthogroups.** Numbers indicate local posterior support values. This topology correctly resolves the shallow relationships that were unresolved in S10 and is otherwise identical to those trees, except for the placement of *E. carneum*.

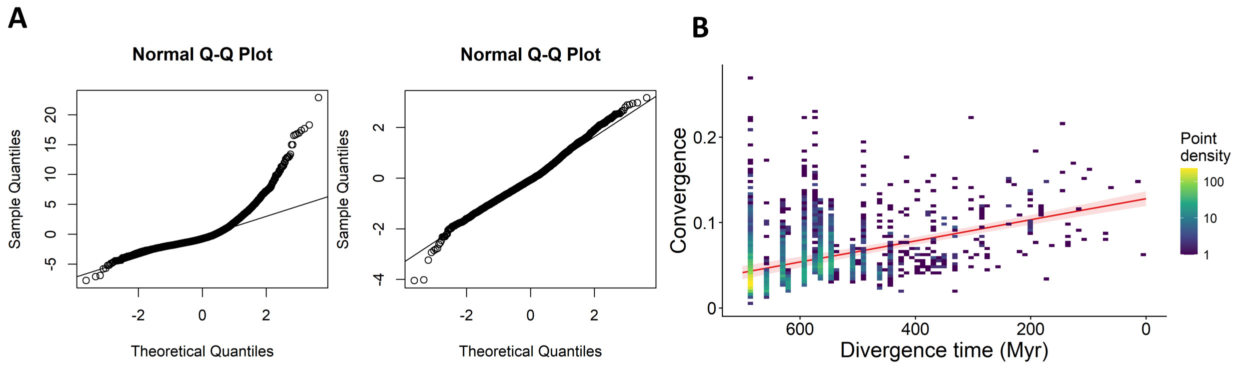


**Figure S12**: **Power transformation improves fit of regression analysis.** A) Quantile-quantile plots showing the residuals of a linear regression of counts against divergence time before (left) and after (right) Yeo-Johnson transformation with $\lambda = -0.565$. B) Phylogenetic regression of untransformed convergence values against divergence time (compare to Fig. 2A.

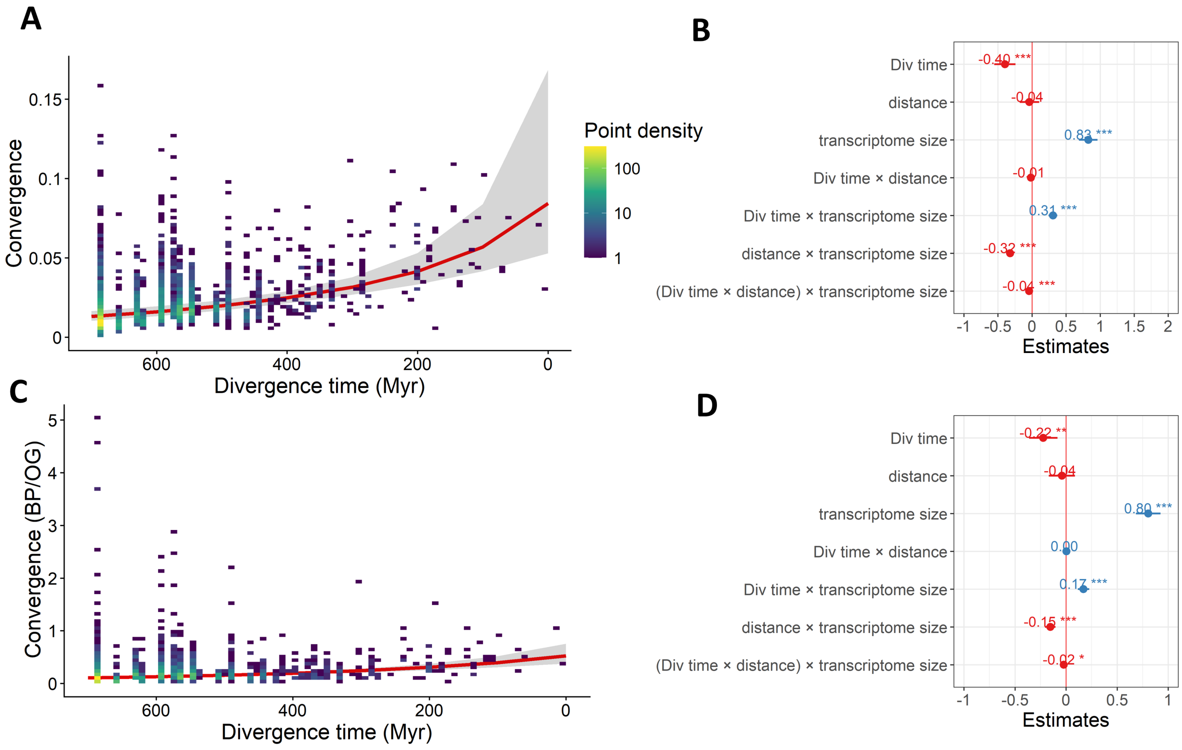


**Figure S13**: **Convergence regression results using a more stringent statistical cutoff and counting the total number of convergence events per OG.** A) Phylogenetic regression of convergence against divergence time using a cutoff of $\omega_c \geq 5$, as opposed to $\omega_c \geq 3$ in the main text. B) Standardized regression coefficients of the regression with all covariates using $\omega_c \geq 5$. C) Phylogenetic regression of convergence against divergence time counting multiple convergence events per OG. The y axis expressed convergence as the total number of convergent branch pairs (BP) over the number of shared OGs. D) Standardized regression coefficients of the regression with all covariates using multiple convergence events per OG. Asterisks indicate significance ($p < 0.05$).

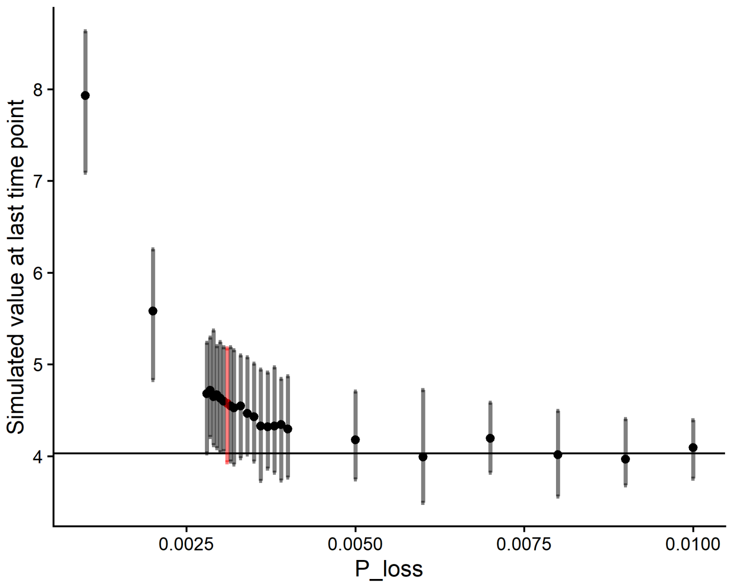


**Figure S14**: **Selection of minimum $P_{loss}$ for null simulations.** Simulated levels of convergence at the oldest time point in our dataset (681.94 Mya) under different values of $P_{loss}$, given $S_0 = 0.3878$ and $S = 0.201$. 500 simulations were performed for each value using two-state Markov chains with vectors of length n=4501. We selected the highest value of $P_{loss}$ whose 95% confidence interval overlapped with our target value, the observed mean level of convergence at 681.94 Mya (solid line; 4.032%). Selected value of $P_{loss}$ (0.0031) is shown in red. For visualization purposes in Fig. 2B, we also analyzed $P_{loss}$ values of 0.0035, 0.005, and 0.01.

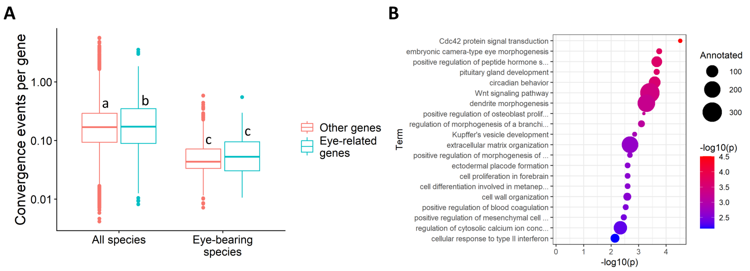


**Figure S15**: **Eye-related genes are only slightly enriched among eye-bearing species.** A) Per-gene convergence rates for eye related (blue, n=827) and non-eye related (red, n=8367) OGs, for all species (left) and comparisons between eye-bearing species (right). Letters indicate significant differences (mixed-effects model followed by Tukey's method). Log scale on y axis. B) Top GO terms enriched among OGs with convergence across 4 independent eye lineages. X axis and colors indicate raw p-values from Fisher's exact test implemented in TopGO. "Annotated" refers to the number of OGs annotated with a corresponding GO term. Full lists of GO terms are in Data 5.

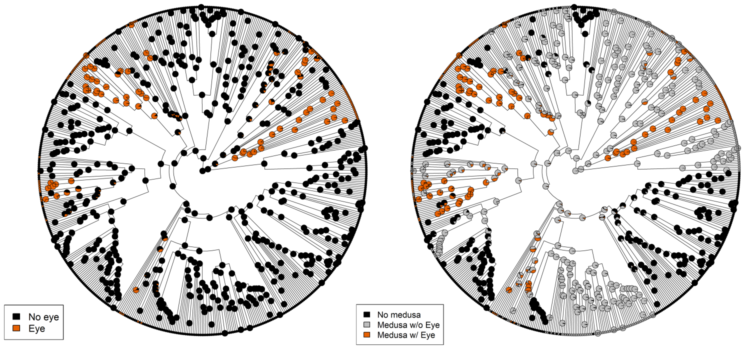


**Figure S16**: **Ancestral state reconstructions (ASR) of eye presence without (left) and with (right) joint modelling of medusa presence.** Orange circles indicate posterior support for eye presence. ASR's were performed using the best-fit models by AIC (Data S4). These analyses inferred at least 12 and 9 eye origins, respectively; the tree on the right is presented in Fig. 1A. Full trees with taxon names can be found on the Github and Dryad.

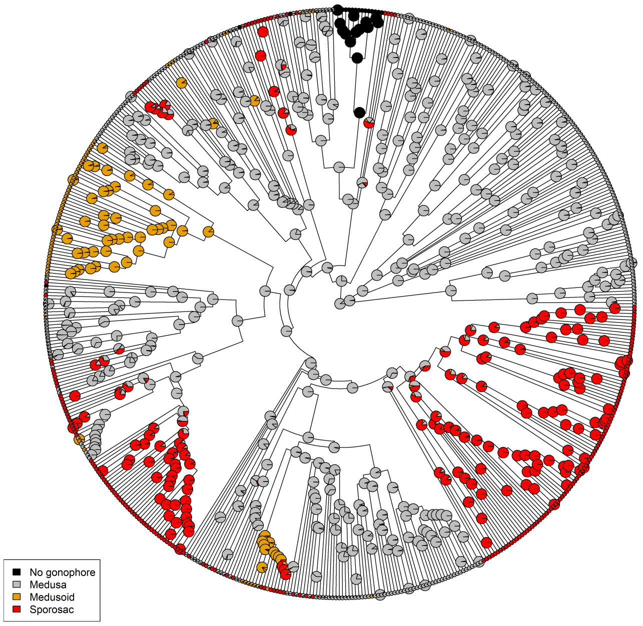


**Figure S17**: **ASR of medusa stage with multi-state encoding.** Circles indicate posterior support for each state. ASR were performed using the best-fit models by AIC (Data S4). The number of inferred medusa losses of all types was identical to that inferred under binary models of medusa presence/absence. Full trees with taxon names can be found on the Github and Dryad.

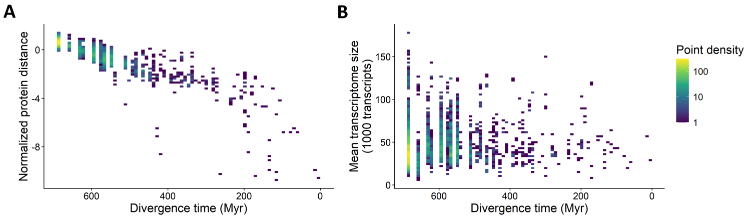


**Figure S18**: **Relationships between median pairwise protein distance (A) and mean transcriptome size (B) with divergence time.**

**Table S1**: **Limited overlapping gene coverage among single-copy orthologs.** Closely-related species with unresolved placements in Fig. S10 have little or no overlap in the single-copy data matrix. This issue is ameliorated by including multi-copy OGs. Data for all species pairs are available on the Dryad.

| Sp1 | Sp2 | shared single-copy orthologs (out of 1742) | shared multi-copy OGs (out of 9194) |
|---|---|---|---|
| Cassiopea andromeda | Cassiopea xamachana | 0 | 3417 |
| Chironex fleckeri | Chironex yamaguchii | 0 | 6518 |
| Haliclystus auricula | Haliclystus octoradiatus | 0 | 5910 |
| Haliclystus auricula | Haliclystus sanjuanensis | 0 | 5535 |
| Hydra sinensis | Hydra viridissima | 0 | 7419 |
| Hydra sinensis | Hydra vulgaris | 0 | 6999 |
| Cyanea capillata | Cyanea nozakii | 1 | 5802 |
| Haliclystus octoradiatus | Haliclystus sanjuanensis | 1 | 5720 |
| Hydra oligactis | Hydra vulgaris | 1 | 6911 |
| Sarsia sp | Sarsia tubulosa | 1 | 5903 |
| Aurelia aurita | Aurelia coerulea | 2 | 7663 |
| Hydra oligactis | Hydra sinensis | 2 | 7065 |
| Hydra viridissima | Hydra vulgaris | 2 | 6970 |
| Millepora alcicornis | Millepora complanata | 2 | 5305 |

**Table S2**: **Convergence in Pax- and Six-family transcription factors.** Pax and Six genes have known roles in cnidarian eye development and experienced convergent substitutions between multiple independent eye lineages. However, they also experienced convergence many times between species without eyes.

| | Orthogroup (OG) | Convergence between eye lineages | Convergence among other species |
|---|---|---|---|
| Pax | OG0000843 | 5 convergent branches across 4 lineages | 91 convergent branches; 74 between 2 lineages w/o eyes |
| Six | OG0001052 | 2 convergent branches across 4 lineages | 113 convergent branches; 94 between 2 lineages w/o eyes |

**Table S3**: **Fits of phylogenetic regression models.** distance: normalized protein distance. tx_size: mean transcriptome size.

| Model | AIC |
|---|---|
| $\sim Div\_time$ | 2637.536 |
| $\sim distance$ | 2660.034 |
| $\sim tx\_size$ | 2545.125 |
| $\sim Div\_time + distance$ | 2638.050 |
| $\sim Div\_time + tx\_size$ | 2468.710 |
| $\sim Div\_time * distance$ | 2632.780 |
| $\sim Div\_time * tx\_size$ | 2431.358 |
| $\sim Div\_time + distance + tx\_size$ | 2456.363 |
| $\sim Div\_time * distance * tx\_size$ | 2219.286 |

**Table S4**: **KEGG terms categorized as "environment-facing" or "housekeeping".**

| KEGG Term | Major category | Categorization |
|---|---|---|
| Xenobiotics | Metabolism | Environment-facing |
| Biosynthesis of other secondary metabolites | Metabolism | Environment-facing |
| Immune System | Organismal Systems | Environment-facing |
| Sensory System | Organismal Systems | Environment-facing |
| Environmental Information Processing | n/a | Environment-facing |
| Cellular Processes | n/a | Housekeeping |
| Nucleotide Metabolism | Metabolism | Housekeeping |
| Genetic Information Processing | n/a | Housekeeping |

**Table S5**: **Fits of phylogenetic regression models.** distance: normalized protein distance. tx_size: mean transcriptome size.

| Model | AIC |
|---|---|
| $\sim Div\_time$ | 2637.536 |
| $\sim distance$ | 2660.034 |
| $\sim tx\_size$ | 2545.125 |
| $\sim Div\_time + distance$ | 2638.050 |
| $\sim Div\_time + tx\_size$ | 2468.710 |
| $\sim Div\_time * distance$ | 2632.780 |
| $\sim Div\_time * tx\_size$ | 2431.358 |
| $\sim Div\_time + distance + tx\_size$ | 2456.363 |
| $\sim Div\_time * distance * tx\_size$ | 2219.286 |

**Table S6**: **Comparisons of independent and correlated models of trait evolution.** Cells contain AIC values with the best-fit model for each combination of characters. There were significant correlations between eyes and medusae and between coloniality and medusae, but not between eyes and coloniality. Medusae were encoded as binary presence/absence.

| | Eye+Medusa | Eyes+Coloniality | Coloniality+Medusa |
|---|---|---|---|
| Independent 1-rate | 523.76 | 583.83 | 740.70 |
| Independent 2-rate | 432.65 | 567.29 | 690.00 |
| Correlated 1-rate | 467.64 | 575.02 | 719.28 |
| Correlated 2-rate | 410.49 | 593.23 | 677.97 |

**Caption for Data S1. Information for transcriptome and genome assemblies.** Accessions, BUSCO scores, and sizes of all assemblies. Unless noted in this file, transcriptomes were assembled from raw reads. All assemblies (peptides and nucleotides) can be found on Dryad.

**Caption for Data S2. Gene ranking for divergence dating analyses by kinda-date.** Genes were ranked according to the 'Weighted_Sum_Ranks' column (lower is better).

**Caption for Data S3. Justifications of fossil calibrations.**

**Caption for Data S4. Ancestral state reconstruction character date and model selection.** Sheets 1 contains character data for medusae and eyes. Where noted, eye references are taken from the supplemental material of (*14*); [SXX] in that columns refers to the relevant citation from that original publication. Sheet 2 contains character data for colonies. Sheet 3 includes details and AIC scores of the various character models tested with corHMM.

**Caption for Data S5. Gene ontology and KEGG enrichment tests.** GO and KEGG annotations can be found on Dryad.
