## Supplementary material for "Adaptive molecular convergence is pervasive across deep time and largely decoupled from phenotypic convergence": Data S3

**Fossil calibrations used in Berger et al.**

**Node:** Cnidaria

**Calibration:** Skew-normal distribution with 563.7 Mya hard minimum and 609 Mya 5% soft maximum ( 'SN(5.637, 0.2312, 183.45)' ).

**Minimum Age:** 563.7 Mya

**Fossil & Justification:** *Haootia quadriformis* from the lower Fermeuse Formation, dated to between 563.67–569.01 Mya (Matthews et al., 2021). Has been interpreted as a stem staurozoan, but this is uncertain (Van Iten et al., 2016). We conservatively use this fossil to constrain crown Cnidaria.

**Maximum Age:** 609 Mya

**Justification:** Age of the Lantian biota, recently dated to 602 +- 7 Mya (Yang et al., 2022). This formation preserves many exceptional algal macrofossils but no definitive crown metazoans.

**Node:** Acraspeda

**Calibration:** 'L(5.419, .1, 0.25, 0.025)'

**Min Age:** 541.9 Mya

**Fossil & Justification:** *Paraconularia ediacara* from the Ediacaran Tamengo Formation (Leme et al. 2022), dated to 542.27 ± 0.38 Mya (Parry et al., 2017). Conulariids may have affinity to scyphozoans or staurozoans, but in any case can reliably constrain crown Acraspeda.

**Node:** Carybdeida

**Calibration:** 'L(5.05, .1, 0.25, 0.025)'

**Min Age:** 505 Mya

**Fossil & Justification:** Specimens UU07021.01 and UU07021.02 from Cartwright et al., 2007. Cambrian Series 3 Marjum Formation (Utah, USA; c. 505 Ma) yields fossils identified as crown cubozoans with likely affinity for Carybdeida (Cartwright et al., 2007; Van Iten et al., 2014).

**Node:** Chirodropidae

**Calibration:** 'L(3.09, .1, 0.25, 0.025)'

**Min Age:** 309 Mya

**Fossil & Justification:** *Anthracomedusa turnbulli,* Upper Carboniferous, Mazon Creek Formation (Young & Hagadorn 2010).

**Node:** Scyphozoa+Cubozoa

**Calibration:** 'L(5.145, .1, 0.25, 0.025)'

**Min Age:** 514.5 Mya

**Fossil & Justification:** *Yunnanoascus haikouensis* from the Lower Cambrian Heilinpu Formation (Han et al., 2016), Cambrian Stage 3 (514.5 - 521 Ma). Explicit phylogenetic analyses place as a stem scyphozoan, so we use this fossil to constrain the split between Scyphozoa and Cubozoa.

**Node:** Rhizostomeae

**Calibration:** 'L(3.05, .1, 0.25, 0.025)'

**Min Age:** 305 Mya

**Fossil & Justification:** *Prothysanostoma eleanorae*, Upper Carboniferous; Cherryvale Formation/Wea Shale, IA. Missourian Stage (Young & Hagadorn 2010).

**Node:** Coronatae

**Calibration:** 'L(5.05, .1, 0.25, 0.025)'

**Min Age:** 505 Mya

**Fossil & Justification:** Specimen UU07021.05 from Cartwright et al., 2007. Cambrian Series 3 Marjum Formation (Utah, USA; c. 505 Ma) yields fossils identified as crown scyphozoans with likely affinity for Coronatae (Cartwright et al., 2007; Van Iten et al., 2014).

**Node:** Hydrozoa

**Calibration:** 'L(5.05, .1, 0.25, 0.025)'

**Min Age:** 505 Mya

**Fossil & Justification:** Cartwright et al, 2007; Van Iten et al., 2014: “The oldest known hydrozoans and cubozoans are represented by medusae in the Cambrian (Series 3) Marjum Formation (Utah, USA; *c*. 505 Ma)”

**Node:** Limnomedusae

**Calibration:** 'L(2.42, .1, 1, 0.025)'

**Min Age:** 242 Mya

**Fossil & Justification:** *Progonionemus vogesiacus*, Lower Triassic; Grès à Voltzia Formation, Middle Triassic (Young & Hagadorn 2010).

**Node:** Plumularioidea

**Calibration:** 'L(3.827, .1, 0.25, 0.025)'

**Min Age:** 382.7 Mya

**Fossil & Justification:** *Archaeoantennularia byersi* Decker, 1952 and

*Plumalina densa* Hall, 1878. Middle Devonian, Sylvania, Michigan, USA and Chemung, Belvidere, New York, USA (Song et al., 2021).

**Node:** Hydractiniidae

**Calibration:** 'L(0.66, .1, 1, 0.025)'

**Min Age:** 66 Mya

**Fossil & Justification:** *Psammoactinia antarctica* Olivero and Aguirre-Urreta, 1994. Late Cretaceous, Early Maastrichtian, Sanctuary Cliffs, Snow Hill Island, Antarctica (Song et al., 2021).

**Node:** Porpitidae (stem)

**Calibration:** 'L(4.211, .1, 0.25, 0.025)'

**Min Age:** 421.1 Mya

**Fossil & Justification:** *Pseudodiscophyllum windermerensis*. Late Silurian, Ludlow, Ludfordian, Bannisdale Formation, England (Fryer and Stanley, 2004).

**Node:** Filifera 1+2 (Proboscidactyla)

**Calibration:** 'L(1.74, .1,1, 0.025)'

**Min Age:** 174 Mya

**Fossil & Justification:** *Protulophila gestroi*. Polish Jura, Lower Jurassic (Słowiński et al., 2020).

**Node:** Scleractinia

**Calibration:** 'L(2.37, .1, 1, 0.025)'

**Min Age:** 237 Mya

**Fossil & Justification:** Many scleractinian fossils appear abruptly in the fossil record at this time (Stanley Jr., 2003).

**Node:** Actiniidae

**Calibration:** 'L(3.09, .1, 0.25, 0.025)'

**Min Age:** 309 Mya

**Fossil & Justification:** *Essexella asherae*. Upper Carboniferous, Mazon Creek Formation (Plotnick et al., 2023).

**Node:** Antipatharia

**Calibration:** 'L(4.70, .1, 0.25, 0.025)'

**Min Age:** 470 Mya

**Fossil & Justification:** *Sinopathes reptans.* Tianjialing, Yichang, Hubei Province, early Arenigian (Baliński et al., 2012)

**Node:** Octocorallia

**Calibration:** 'L(1.36, .1, 1, 0.025)'

**Min Age:** 136 Mya

**Fossil & Justification:** *Pseudopolytremacis japonica*. Belvédère de Gaud, SW Saint-Remèze, Ardeche, Upper Barremian (Löser & Ferry, 2006).

**Node:** Cnidaria

**Calibration:** Skew-normal distribution with 563.7 Mya hard minimum and 609 Mya 5% soft maximum ( 'SN(5.637, 0.2312, 183.45)' ).

**Minimum Age:** 563.7 Mya

**Fossil & Justification:** *Haootia quadriformis* from the lower Fermeuse Formation, dated to between 563.67–569.01 Mya (Matthews et al., 2021). Has been interpreted as a stem staurozoan, but this is uncertain (Van Iten et al., 2016). We conservatively use this fossil to constrain crown Cnidaria.

**Maximum Age:** 609 Mya

**Justification:** Age of the Lantian biota, recently dated to 602 +- 7 Mya (Yang et al., 2022). This formation preserves many exceptional algal macrofossils but no definitive crown metazoans.

**Node:** Acraspeda

**Calibration:** 'L(5.419, .1, 0.25, 0.025)'

**Min Age:** 541.9 Mya

**Fossil & Justification:** *Paraconularia ediacara* from the Ediacaran Tamengo Formation (Leme et al. 2022), dated to 542.27 ± 0.38 Mya (Parry et al., 2017). Conulariids may have affinity to scyphozoans or staurozoans, but in any case can reliably constrain crown Acraspeda.

**Node:** Carybdeida

**Calibration:** 'L(5.05, .1, 0.25, 0.025)'

**Min Age:** 505 Mya

**Fossil & Justification:** Specimens UU07021.01 and UU07021.02 from Cartwright et al., 2007. Cambrian Series 3 Marjum Formation (Utah, USA; c. 505 Ma) yields fossils identified as crown cubozoans with likely affinity for Carybdeida (Cartwright et al., 2007; Van Iten et al., 2014).

**Node:** Chirodropidae

**Calibration:** 'L(3.09, .1, 0.25, 0.025)'

**Min Age:** 309 Mya

**Fossil & Justification:** *Anthracomedusa turnbulli,* Upper Carboniferous, Mazon Creek Formation (Young & Hagadorn 2010).

**Node:** Scyphozoa+Cubozoa

**Calibration:** 'L(5.145, .1, 0.25, 0.025)'

**Min Age:** 514.5 Mya

**Fossil & Justification:** *Yunnanoascus haikouensis* from the Lower Cambrian Heilinpu Formation (Han et al., 2016), Cambrian Stage 3 (514.5 - 521 Ma). Explicit phylogenetic analyses place as a stem scyphozoan, so we use this fossil to constrain the split between Scyphozoa and Cubozoa.

**Node:** Rhizostomeae

**Calibration:** 'L(3.05, .1, 0.25, 0.025)'

**Min Age:** 305 Mya

**Fossil & Justification:** *Prothysanostoma eleanorae*, Upper Carboniferous; Cherryvale Formation/Wea Shale, IA. Missourian Stage (Young & Hagadorn 2010).

**Node:** Coronatae

**Calibration:** 'L(5.05, .1, 0.25, 0.025)'

**Min Age:** 505 Mya

**Fossil & Justification:** Specimen UU07021.05 from Cartwright et al., 2007. Cambrian Series 3 Marjum Formation (Utah, USA; c. 505 Ma) yields fossils identified as crown scyphozoans with likely affinity for Coronatae (Cartwright et al., 2007; Van Iten et al., 2014).

**Node:** Hydrozoa

**Calibration:** 'L(5.05, .1, 0.25, 0.025)'

**Min Age:** 505 Mya

**Fossil & Justification:** Cartwright et al, 2007; Van Iten et al., 2014: “The oldest known hydrozoans and cubozoans are represented by medusae in the Cambrian (Series 3) Marjum Formation (Utah, USA; *c*. 505 Ma)”

**Node:** Limnomedusae

**Calibration:** 'L(2.42, .1, 1, 0.025)'

**Min Age:** 242 Mya

**Fossil & Justification:** *Progonionemus vogesiacus*, Lower Triassic; Grès à Voltzia Formation, Middle Triassic (Young & Hagadorn 2010).

**Node:** Plumularioidea

**Calibration:** 'L(3.827, .1, 0.25, 0.025)'

**Min Age:** 382.7 Mya

**Fossil & Justification:** *Archaeoantennularia byersi* Decker, 1952 and

*Plumalina densa* Hall, 1878. Middle Devonian, Sylvania, Michigan, USA and Chemung, Belvidere, New York, USA (Song et al., 2021).

**Node:** Hydractiniidae

**Calibration:** 'L(0.66, .1, 1, 0.025)'

**Min Age:** 66 Mya

**Fossil & Justification:** *Psammoactinia antarctica* Olivero and Aguirre-Urreta, 1994. Late Cretaceous, Early Maastrichtian, Sanctuary Cliffs, Snow Hill Island, Antarctica (Song et al., 2021).

**Node:** Porpitidae (stem)

**Calibration:** 'L(4.211, .1, 0.25, 0.025)'

**Min Age:** 421.1 Mya

**Fossil & Justification:** *Pseudodiscophyllum windermerensis*. Late Silurian, Ludlow, Ludfordian, Bannisdale Formation, England (Fryer and Stanley, 2004).

**Node:** Filifera 1+2 (Proboscidactyla)

**Calibration:** 'L(1.74, .1,1, 0.025)'

**Min Age:** 174 Mya

**Fossil & Justification:** *Protulophila gestroi*. Polish Jura, Lower Jurassic (Słowiński et al., 2020).

**Node:** Scleractinia

**Calibration:** 'L(2.37, .1, 1, 0.025)'

**Min Age:** 237 Mya

**Fossil & Justification:** Many scleractinian fossils appear abruptly in the fossil record at this time (Stanley Jr., 2003).

**Node:** Actiniidae

**Calibration:** 'L(3.09, .1, 0.25, 0.025)'

**Min Age:** 309 Mya

**Fossil & Justification:** *Essexella asherae*. Upper Carboniferous, Mazon Creek Formation (Plotnick et al., 2023).

**Node:** Antipatharia

**Calibration:** 'L(4.70, .1, 0.25, 0.025)'

**Min Age:** 470 Mya

**Fossil & Justification:** *Sinopathes reptans.* Tianjialing, Yichang, Hubei Province, early Arenigian (Baliński et al., 2012)

**Node:** Octocorallia

**Calibration:** 'L(1.36, .1, 1, 0.025)'

**Min Age:** 136 Mya

**Fossil & Justification:** *Pseudopolytremacis japonica*. Belvédère de Gaud, SW Saint-Remèze, Ardeche, Upper Barremian (Löser & Ferry, 2006).
